# Stromal Prostaglandin is a Dominant Spatial Regulator of Cell-fate Plasticity in Colorectal Cancer

**DOI:** 10.64898/2026.06.14.732116

**Authors:** Corinne Molyneux, Rhianna O’Sullivan, Eoghan J. Mulholland-Illingworth, Joshua W. Moore, Nick Li, Petra Vlckova, Raheleh Amirkhah, Aurélie Dobric, Shauna Crampsie, Alistair Wilkinson, Jack Langley, Marta Labarquilla Alonso, Ashley Campbell, Jeroen Claus, Smita Krishnaswamy, Philip D. Dunne, Simon Leedham, Christopher J. Tape

## Abstract

Colorectal cancer (CRC) tumours with high stromal content have a worse outcome, but the mechanisms governing this are unclear. Using high-throughput single-cell perturbation analysis of CRC patient-derived organoids (PDOs) and cancer-associated fibroblasts (CAFs) we find that epithelial cells with high stromal-communication potential are marked by the transcriptional co-repressor *DACH1*. To define the causal regulators of stromal-epithelial signalling, we developed a novel CRISPR screening platform to perturb the CAF secretome and measure epithelial stem cell responses at single-cell resolution. Intercellular CRISPR screening and full factorial ligand analysis revealed that stromal Prostaglandin E2 (PGE_2_) is a dominant regulator of CRC cell-fate plasticity. Stromal PGE_2_ converts *DACH1*^*+*^ epithelia from a chemosensitive proliferative colonic stem cell (proCSC) fate into a chemorefractory and prometastatic revival colonic stem cell (revCSC) fate. PGE_2_-driven epithelial transdetermination is rapid and reversible, providing an acute mechanism for stromal-driven plasticity in CRC tumours. Genetic and pharmacological inhibition of stromal COX2 inhibits epithelial plasticity, trapping CRC epithelia in an anti-metastatic and chemosensitive proCSC fate. *PTGS2*^*+*^ CAFs support a spatially resolved revCSC to proCSC plasticity gradient in human CRC tumours marked by increasing *DACH1* expression. These results reveal that stromal prostaglandin is a dominant spatial regulator of poor-prognosis cell-fates and may explain the benefit of anti-COX therapies in both preventing and treating CRC.

## Introduction

Colorectal cancer (CRC) afflicts >1.9 million people a year worldwide and is a major source of cancer-related deaths [1]. CRC tumours are heterocellular systems that comprise mutated epithelial cancer cells, stromal cancer-associated fibroblasts (CAFs), and various immune cells [2]. Once considered a purely genetically-driven disease, it is now widely appreciated that CRC is regulated by both cell-intrinsic mutations and non-genetic cell-extrinsic cues from the tumour microenvironment [3, 4]. Epithelial CRC cells are phenotypically plastic and occupy a range of cell-fates spanning from proliferative colonic stem cells (proCSC), revival colonic stem cells (revCSC) (also known as onco-foetal cells), and multiple non-canonical secretory and absorptive lineages [5, 3, 6]. proCSC express canonical intestinal stem cell genes (*LGR5*^*+*^, *OLFM4*^*+*^) but are hyperproliferative relative to untransformed colonic stem cells (CSCs). By contrast, revCSC are typically slower-cycling and express a foetal-like gene signature (*ANXA1*^*+*^, *PKN2*^*+*^, and *L1CAM*^*+*^). While proCSC are responsible for tumour growth, revCSC are often found at the invasive front of CRC tumours [7], play an essential role in metastasis [8], and provide a developmental gateway to non-canonical differentiation in advanced disease [9]. This non-genetic plasticity enables CRC tumours to expand when unchallenged and survive when stressed [10].

CRC tumours with a high abundance of CAFs (originally identified as the consensus molecular subtype (CMS) 4 [11]) typically have increased metastasis, poor response to chemotherapy, and a worse prognosis [12, 13]. We recently showed that CAFs can transdeterminate proCSC cancer cells towards a revCSC-like fate – providing evidence that stromal-driven epithelial plasticity can regulate chemosensitivity [14]. However, we also found that cell-intrinsic mutations change how epithelial cells respond to stromal cues [3] and that not all patient-derived organoids (PDOs) effectively communicate with CAFs [14]. Dozens of CAF-epithelial communication ligands have been identified in CRC [2, 10], but the precise intercellular signalling factors responsible for CAF-induced epithelial plasticity in human tumours are undefined.

Despite the clinical importance of epithelial plasticity and stromal CAFs in CRC, it is currently unclear exactly how CAFs regulate epithelial cell-fate and which epithelial cancer cells can undergo plasticity shifts. Here, using systematic perturbation of CRC PDOs and CAFs and high-throughput multimodal single-cell analysis, we functionally define how CAFs regulate CRC plasticity. We find that stromal-plasticity regulation is dependent on the cell-intrinsic state of CRC epithelia. Specifically, we find that *DACH1*^*+*^ cancer cells are highly responsive to CAFs, whereas *DACH1*^*–*^ cancer cells are not regulated by CAFs. To define the native CAF cues that regulate epithelial plasticity we developed a novel intercellular signalling CRISPR screening platform. Single-cell CAF-PDO secretome CRISPR screening revealed that CAF-derived IL1R1, HGF, GREM1, and PGE_2_ can regulate proCSC to revCSC transdetermination, but PGE_2_ is a master regulator of CRC plasticity. Stromal PGE_2_ signals via MAPK, PI3K, and AP-1 to drive revCSC and both genetic and pharmacological inhibition of stromal COX2 blocks epithelial plasticity. Finally, we find that *PTGS2*^*+*^ CAFs form a localised epithelial plasticity gradient that is mapped by *DACH1* in human CRC tumours. These results demonstrate that prostaglandin is a dominant stromal regulator of CRC plasticity and provide a mechanistic rationale for anti-PGE_2_ therapies in CRC.

## Results

### Stromal Regulation of CRC Plasticity

CAFs can regulate CRC epithelia, but the intercellular mechanisms governing this process are unclear. To investigate the influence of CAFs across a range of CRC cells, we first performed scRNA-seq analysis of x10 CRC PDOs co-cultured +/-CRC CAFs in triplicate (x90 experimental conditions, 89,894 PDOs, 71,004 CAFs). Waddington-like Valley Ridge (VR) landscape analysis [3] revealed that CRC PDOs comprise an admixture of cells found in CRC tumours *in vivo*, notably mitotic proCSC (*RRM2*^+^, *HELLS*^+^, *TOP2A*^+^, *MKI67* ^+^) and slow-cycling revCSC (*CD55*^+^, *EMP1*^+^, *CLDN4*^+^, *ANXA1*^+^), with additional non-canonically differentiated neuroendocrine and squamous-like cells (Figure 1a, Figure S1a-f). The proCSC-revCSC axis can be clearly observed using a colonic stem cell index (SCI) [5] across all PDOs (Figure S1g), with each PDO displaying a patient-specific baseline stem cell admixture. In parallel to scRNA-seq analysis, we also determined the chemosensitivity of all CRC PDOs and found that intrinsic stem cell admixture significantly correlates with chemosensitivity (Figure 1b). Specifically, the relative proCSC score of each PDO significantly correlated with chemosensitivity to 5-FU, SN-38 (irinotecan), and oxaliplatin. Transcription factor activity analysis [15] further revealed cell-type specific transcriptional programmes across all PDOs (Figure 1c). We found that proCSC have high E2F1/4, ZEB1, TWIST2, SNAI2, ASCL2, and SOX9 transcription factor activity, whereas revCSC have high TEAD1, JUND, AP-1, TCF7, RXRB, STAT6, and SMAD6 activity. Differentiated cells also displayed unique transcription factor activity, with neuroendocrine cells employing NEUROG3, squamous cells activating IKZF4, and goblet-like cells demonstrating high GATA5 activity. 5-FU, irinotecan, and oxaliplatin each target different molecular processes and we found that distinct proCSC and revCSC transcription factor activities significantly correlate with specific chemotherapy responses across all CRC PDOs. For example, 5-FU sensitivity strongly aligns with SNAI2^High^ and SMAD6^Low^, irinotecan with E2F4^High^ and RXRB^Low^, and oxaliplatin with SOX9^High^ and SMAD6^Low^ transcription factor activity (Figure S1h). These results underscore the importance of epithelial cell-fate in controlling response to standard-of-care CRC chemotherapies.

**Figure 1.**
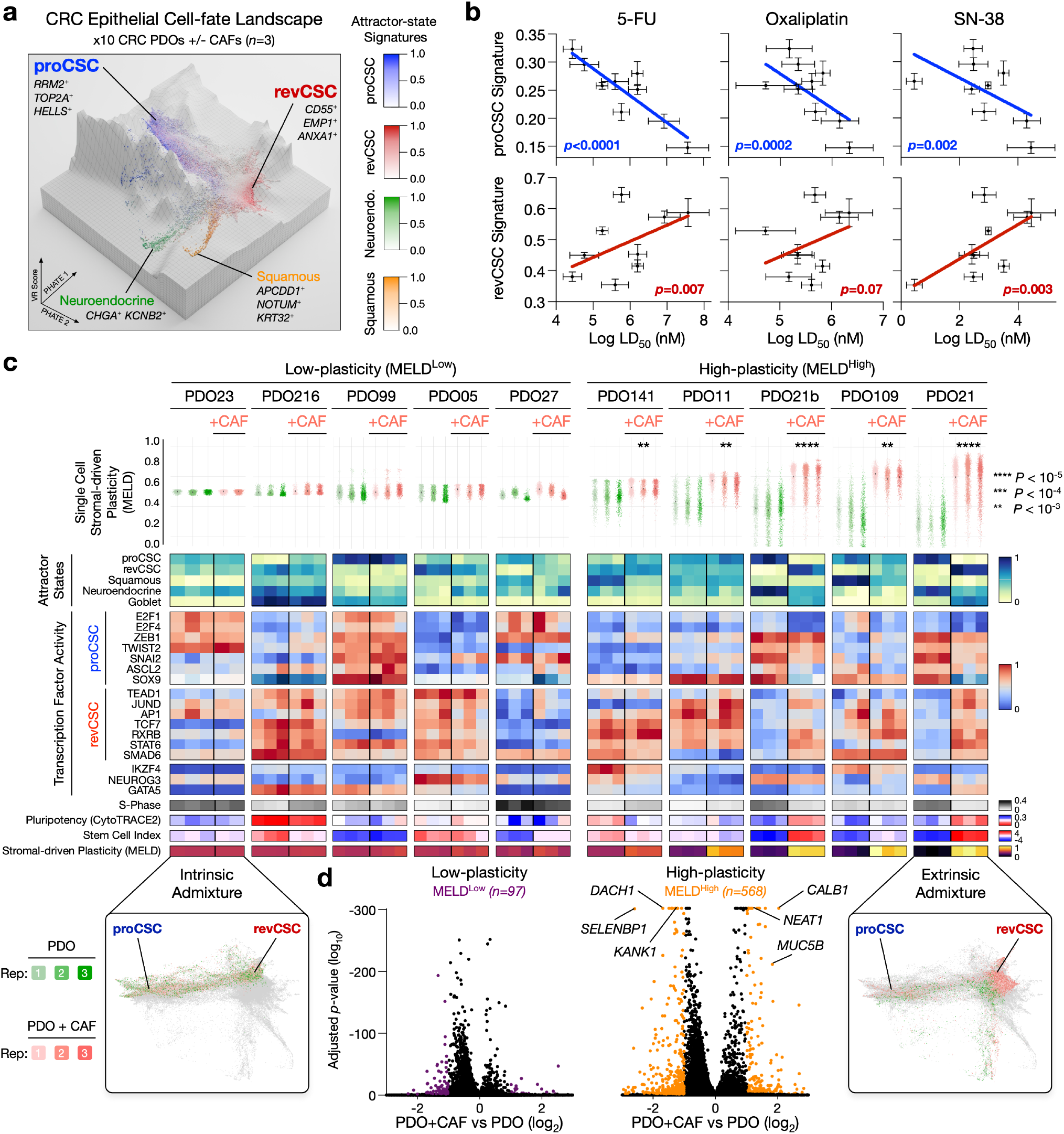
CRC Epithelial State Predicts Chemosensitivity and Stromal Plasticity Response. **a)** scRNA-seq VR landscape of x10 CRC PDOs +/-CAFs (in triplicate) coloured by gene signatures to highlight plasticity attractor states (89,894 single cells). **b)** PDO proCSC and revCSC gene signatures vs 5-FU, Oxaliplatin, and SN-38 (Irinotecan) LD_50_. Pearson correlation. **c)** Single-cell transcriptional features of PDOs +/-CAFs. **d)** Differential gene expression regulation by CAFs in MELD^Low^ and MELD^High^ PDOs.

To determine how CRC plasticity is regulated by CAFs, we computed the MELD-likelihood score [16] per PDO +/-CAFs as a single-cell metric of CAF-induced epithelial transcriptional change and analysed downstream transcription factor activity, gene signatures, cell-cycle activity, putative pluripotency, and stem cell index (Figure 1c). MELD allowed us to directly quantify how each epithelial population transcriptionally responds to CAFs – providing a statistical measurement of stromal-induced plasticity per PDO. For example, PDO23 comprises an intrinsic proCSC/revCSC stem cell admixture that is not significantly regulated by CAFs (low-plasticity). Conversely, PDO21 is intrinsically proCSC, but significantly transdeterminates to revCSC when communicating with CAFs (high-plasticity). Using MELD significance across replicates, we assigned each PDO into ‘low-plasticity’ (MELD^Low^) and ‘high-plasticity’ (MELD^High^) groups to explore common features of stromal-reactive cancer cells. As expected, CAFs significantly regulate more genes in high-plasticity MELD^High^ PDOs (*n*=568) than low-plasticity MELD^Low^ PDOs (*n*=97), including upregulation of *CALB1, NEAT1*, and various mucins (*MUC2, MUC5B*). Interestingly, CAFs consistently downregulate the Hippo pathway regulator *KANK1* and the AP-1 transcriptional co-repressor *DACH1* (Dachshund homolog 1) (Figure 1d) in high-plasticity PDOs. CAFs regulate a range of transcription factor activity in MELD^High^ PDOs, including a downregulation of the canonical intestinal identity markers CDX2 and SOX9, and upregulation of hypoxia (HIF1A, EPAS1, ATF4) and inflammatory (STAT1/2, RELA, NFKB) activity (Figure S1i). Collectively, these results suggest that CRC cells have cell-intrinsic identities that dictate response to chemotherapy and patient-specific plasticity responses to stromal cues.

### CAF-driven Plasticity is Dictated by *DACH1*^*+*^ Cancer Cells

We next sought to define what dictates CAF-induced plasticity: CAF cell-fate, CAF intercellular signalling, or epithelial cell-fate. While CRC CAFs adopt a myofibroblast-like state in monoculture, we found that all x10 CRC PDOs polarise CAFs towards a ‘tumour-like’ CAF transcriptional subtype [17] in 3D co-culture (Figure 2a, Figure S2a-b). However, this tumour-like CAF gene signature is induced by both MELD^High^ (high-plasticity) and MELD^Low^ (low-plasticity) PDOs and therefore cannot explain why some PDOs respond to CAFs and others do not (Figure 2b). We next found that PDOs regulate the intercellular communication potential of CAFs, including upregulation of *VEGFA/C* and the inflammatory prostaglandin synthase gene *PTGS2* (Figure 2c). However, there are minimal differences between CAF intercel-lular communication genes in MELD^High^ and MELD^Low^ PDOs (Figure 2d), suggesting that CAF intercellular ligands are necessary but not sufficient to drive CRC epithelial plasticity.

**Figure 2.**
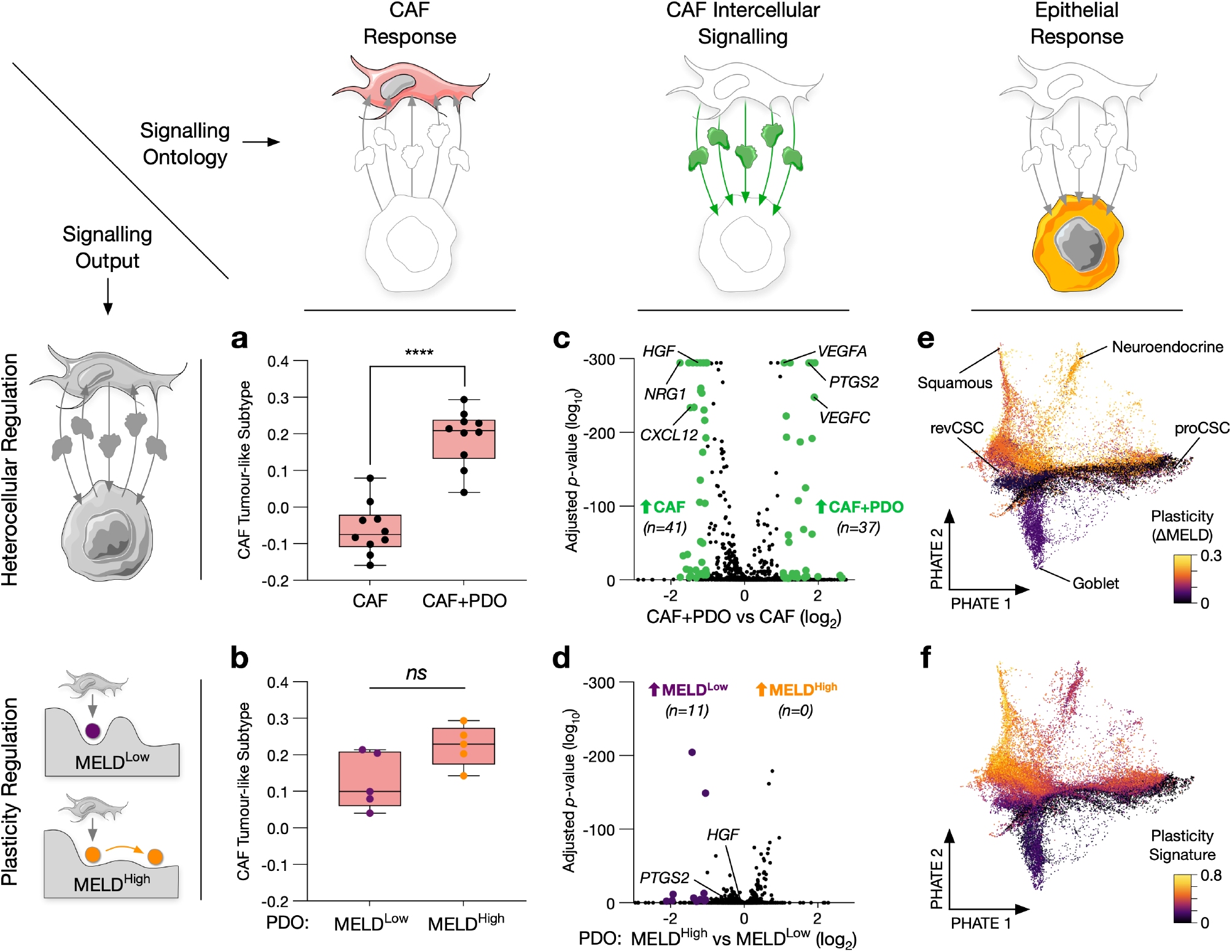
CAF-driven Plasticity Responses are Dictated by CRC Epithelial Cells. **a)** Tumour-like CAF gene signature in CAFs +/-x10 different PDOs. **b)** Tumour-like CAF gene signature in CAFs co-cultured with MELD^High^ and MELD^Low^ PDOs. **c)** Regulation of CAF intercellular signalling genes +/-PDOs. **d)** Regulation of CAF intercellular signalling genes in MELD^High^ and MELD^Low^ PDOs. **e)** Single cell PHATE of PDO monoculture (39,022 cells) coloured by CAF-induced plasticity (ΔMELD), or **f)** stromal plasticity gene signature. Paired *t*-test. *** = *p* < 0.001, ** = *p* < 0.01, * = *p* < 0.05, ns= not significant.

To determine why some PDOs are sensitive to stromal communication while others are not, we next compared the baseline transcriptome of CRC epithelia to their CAF-induced plasticity response (ΔMELD). Integrated analysis of scRNA-seq data from x10 CRC PDOs in monoculture (in triplicate, 39,022 single cells) clearly identified proCSC, revCSC, squamous, neuroendocrine, and goblet-like attractor states (Figure S2c-j). Strikingly, we found that CAF-induced plasticity strongly aligns with the cell-intrinsic epithelial transcriptome (Figure 2e). Epithelial plasticity potential did not correlate with stem cell index or any epithelial cell-type (Figure S2k), but could instead be captured by an epithelial plasticity response gene signature that spanned various proCSC, revCSC, neuroendocrine, and squamous cells (Figure 2f). Collectively, these results suggest that in CRC cell-extrinsic plasticity potential is determined by the cell-intrinsic transcriptional state of epithelial cells.

When viewed in a Waddington-like VR landscape [3], stromal-plasticity response does not correlate with predicted epithelial pluripotency (Figure 3a) (Figure S2i). Instead, high-plasticity CRC PDOs typically contain non-canonical squamous and neuroendocrine cells, whereas low-plasticity PDOs are enriched for secretory goblet cells. Specifically, epithelia that can be reprogrammed by CAFs uniformly express the transcriptional co-repressor *DACH1*, as well as *NRXN3* and *MGAM2*, whereas PDOs that cannot communicate with CAFs express *GRB10, DCBLD2*, and *TENM3* (Figure 3b-c). We found that *DACH1* can act as a single-gene marker of stromal plasticity response (Figure 3d-f). In CRC patients [18], *DACH1* does not mark a specific cell-type, but is expressed in proCSC, squamous, and neuroendocrine cells, with peak expression as cells transition along the proCSC to revCSC axis (Figure S2l-n). We find that *DACH1* can clearly resolve high- and low-plasticity potential cancer cells (Figure 3g) and *DACH1* is further downregulated by intercellular signalling from CAFs in high-plasticity PDOs (Figure 3h). Collectively, these results suggest that *DACH1* is a marker of epithelial competence for stromal reprogramming in CRC.

**Figure 3.**
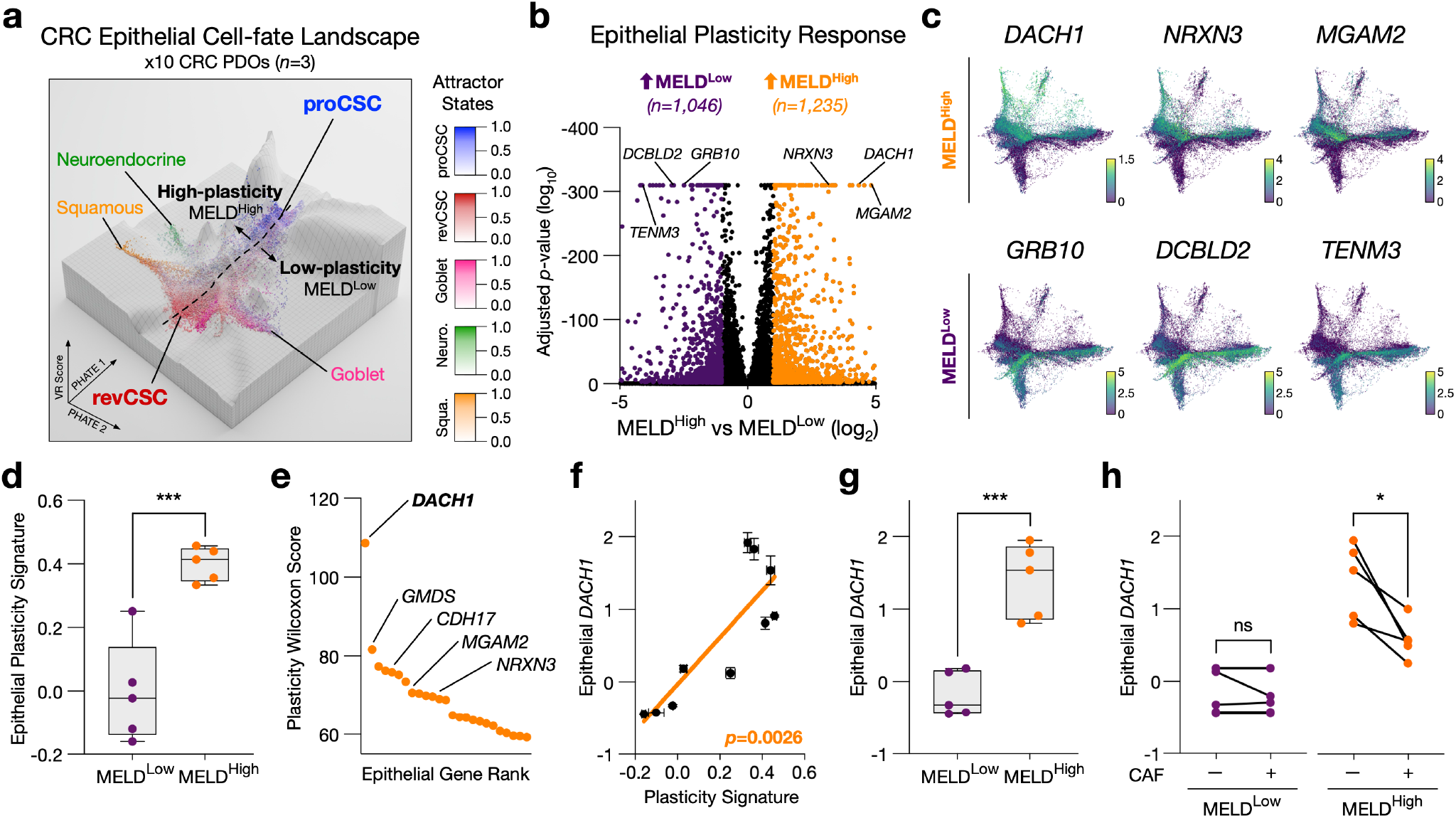
High-plasticity CRC Epithelia are *DACH1*^*+*^. **a)** scRNA-seq VR landscape of x10 CRC PDO monocultures (in triplicate) (39,022 single cells), coloured by CRC attractor state gene signatures. **b)** Regulation of epithelial genes between MELD^High^ (high-plasticity) and MELD^Low^ (low-plasticity) PDOs. **c)** Single-cell PHATE coloured by MELD^High^ and MELD^Low^ genes. **d)** PDO plasticity gene signature per PDO grouped by MELD^High^ and MELD^Low^. Unpaired *t*-test. **e)** Plasticity Wilcoxon score for MELD^High^ genes. **f)** PDO *DACH1* expression versus PDO plasticity signature per PDO. Pearson correlation. **g)** PDO *DACH1* expression per PDO grouped by MELD^High^ and MELD^Low^. Unpaired *t*-test. **h)** PDO *DACH1* expression +/-CAFs per PDO grouped by MELD^High^ and MELD^Low^. Paired *t*-test. *** = *p* < 0.001, ** = *p* < 0.01, * = *p* < 0.05, ns= not significant.

### Single-cell CRISPR Screening of Intercellular Signalling Identifies CAF Stromal Regulators of CRC Epithelial Plasticity

We next sought to identify the intercellular CAF signals that regulate *DACH1*^*+*^ CRC epithelial plasticity. Treating *DACH1*^*+*^ CRC PDOs with CRC CAF-conditioned media phenocopied direct CAF co-culture (Figure S3a-c), confirming that intercellular communication is dependent on soluble ligands from the CAF secretome. While computational ligand-receptor analysis can identify putative intercellular communication between cells, experimental validation is required to determine functional cell-cell signalling ligands. To functionally identify the CAF ligands driving epithelial CRC plasticity, we developed a novel single-cell intercellular signalling CRISPR screening platform (Figure 4a). We first identified the putative CRC CAF secretome using a combination of ligand-receptor analysis of PDO-CAF scRNA-seq data, cytokine array analysis of CAF-conditioned media, and the literature (resulting in x202 CAF ligand targets). We then reverse-transfected Cas9^+^ CRC CAFs (Figure S3d) with gRNAs to individually knock out (KO) CAF ligands (85-99% KO efficiency of native CAF ligands (Figure S3e)) before co-culturing KO CAFs with *DACH1*^*+*^ CRC PDOs in an arrayed format. PDO-CAF KO co-cultures were then analysed by thiol-organoid barcoding *in situ* mass cytometry (TOB*is* MC) [19, 20] – measuring post-translational modification (PTM) signalling, cell-cycle, and cell-fate at single-cell resolution in both PDOs and CAFs for each lig- and KO across x4 biological replicates. This experimental system enabled us to determine how native ligands from the CAF secretome functionally regulate epithelial CRC plasticity.

**Figure 4.**
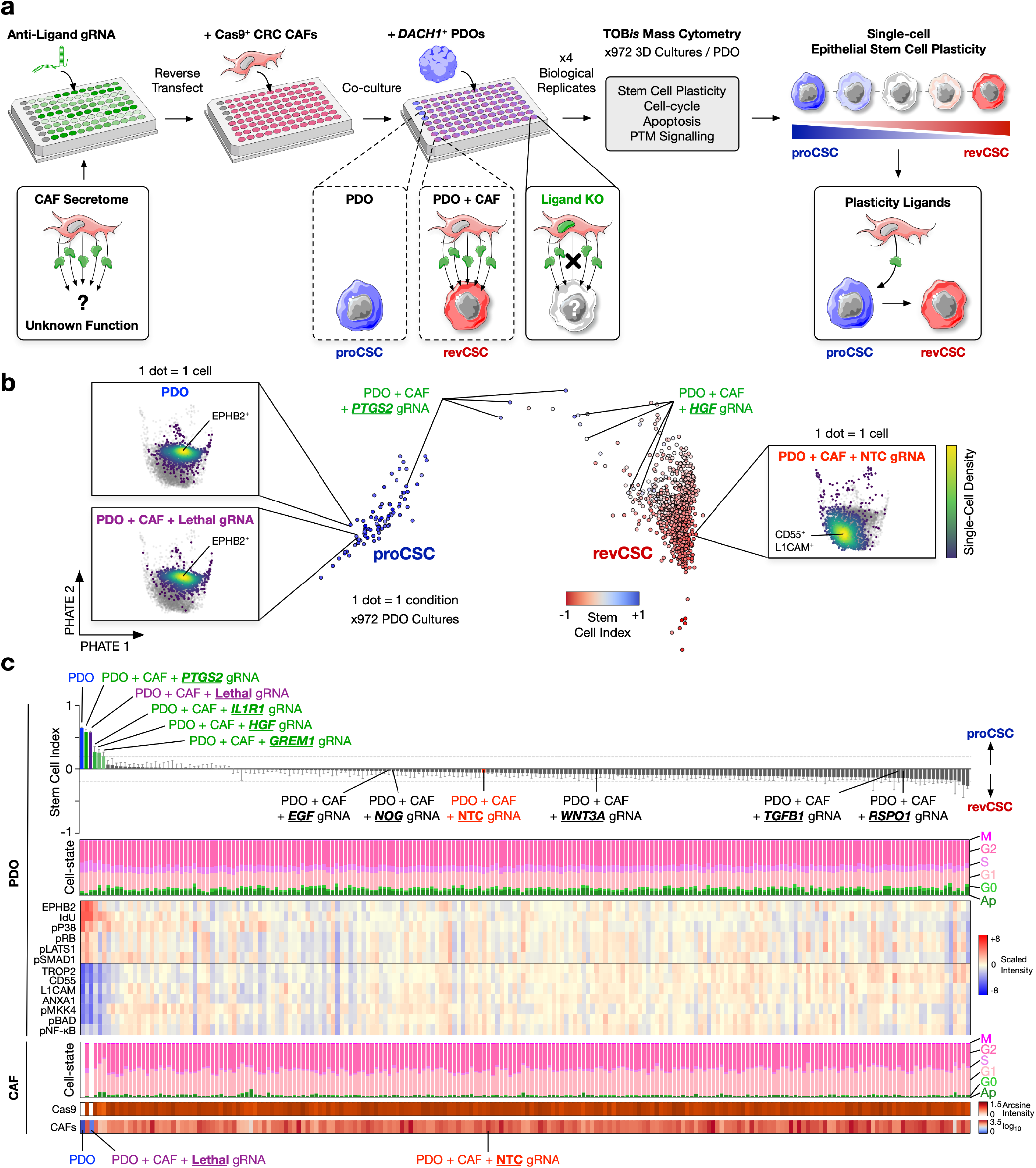
Single-cell Intercellular Signalling CRISPR Screening Reveals CAF *PTGS2, IL1R1, GREM1*, and *HGF* can Regulate Epithelial Plasticity. **a)**. Single-cell intercellular signalling CRISPR screening experimental workflow. **b)** Geodesic sinkhorn PHATE phenoscape of x972 PDO cultures coloured by stem cell index (SCI) (1 dot = 1 condition). Single-cell density PHATE plots of PDO monoculture, PDO + CAF + non-targeting control (NTC) gRNA, and PDO + CAF + lethal gRNA (1 dot = 1 cell). **c)** Stem cell index, cell-state, and EMD protein and signalling for each PDO condition (*n*=4). CAF cell-state, Cas9 expression, and CAF cell numbers per condition (*n*=4).

Single-cell analysis of x972 PDO-CAF KO co-cultures revealed cell-type specific regulation of intercellular signalling. Using Geodesic Sinkhorn optimal transport [21] to develop a condition-level phenoscape embedding, we could clearly observe that PDOs alone retained their intrinsic proCSC-fate, whereas PDOs + Cas9^+^ CAFs + non-targeting control (NTC) gRNA are revCSC-enriched (Figure 4b). Crucially, PDOs + Cas9^+^ CAFs + lethal gRNA remained proCSC and retained <1% CAFs compared to NTC (Figure S3f). We used B7-H3 (CD276) as a non-functional protein knockout control and CAF B7-H3 levels were significantly depleted by anti-B7-H3 gRNA in all replicates (Figure S3g). Collectively, these data confirm high CAF CRISPR editing efficiency throughout the intercellular signalling screening assay.

To quantify the effect of CAF ligand KO on CRC epithelial plasticity, we computed the proCSC-revCSC SCI for each co-culture (Figure 4b, Figure S3h). In agreement with our previous observation that CRC oncogenic mutations can inhibit homeostatic stromal-epithelial communication [3], KO of homeostatic stromal ligands *WNT3A, EGF, NOG*, and *RSPO1* had no effect on CRC PDO SCI (Figure 4c). Despite the well-documented role of stromal TGF-β in murine models of CRC [3], we also found that perturbing native TGF-β in human CAFs had no influence on CRC epithelia. By contrast, CAF KO of *PTGS2, IL1R1, GREM1*, or *HGF* all increased epithelial SCI across x4 biological replicates, enabling PDOs to remain in a proCSC-enriched state while in co-culture with CAFs. Specifically, CRISPR KO of CAF *PTGS2* maintained a similar epithelial SCI to monoculture PDOs, increasing EPHB2 expression and S-phase entry, while decreasing expression of revCSC markers CD55, TROP2, L1CAM, and ANXA1. *PTGS2* KO CAFs also increased PDO proCSC signalling via pP38-MAPK [T180/Y182] and pRB [S807/811], while decreasing revCSC signalling via pNF-κB [S529], pBAD [S112], pMKK4 [S257], and YAP (pLATS1/2 [S872/909]).

### CAF PGE_2_ is a Dominant Regulator of Stromal-induced Plasticity in CRC

To understand how PGE_2_, IL1R1, GREM1, and HGF regulate epithelial stem cell plasticity, we treated CRC PDOs with all combinations of all ligands in a full 2^4^ facto-rial array (in triplicate) and analysed the single-cell PTM signalling, cell-cycle, and cell-fate plasticity responses of each PDO by TOB*is* MC. To confirm these results were not anecdotal to the discovery PDO used in the initial CRISPR screen, we also treated x3 new CRC PDOs with all 2^4^ factorial array ligand combinations. Untreated PDO monocultures provided a proCSC control and PDO+CAF co-cultures acted as an intercellular signalling-driven revCSC control (x51 treatments per PDO, x204 single-cell datasets) (Figure 5a).

**Figure 5.**
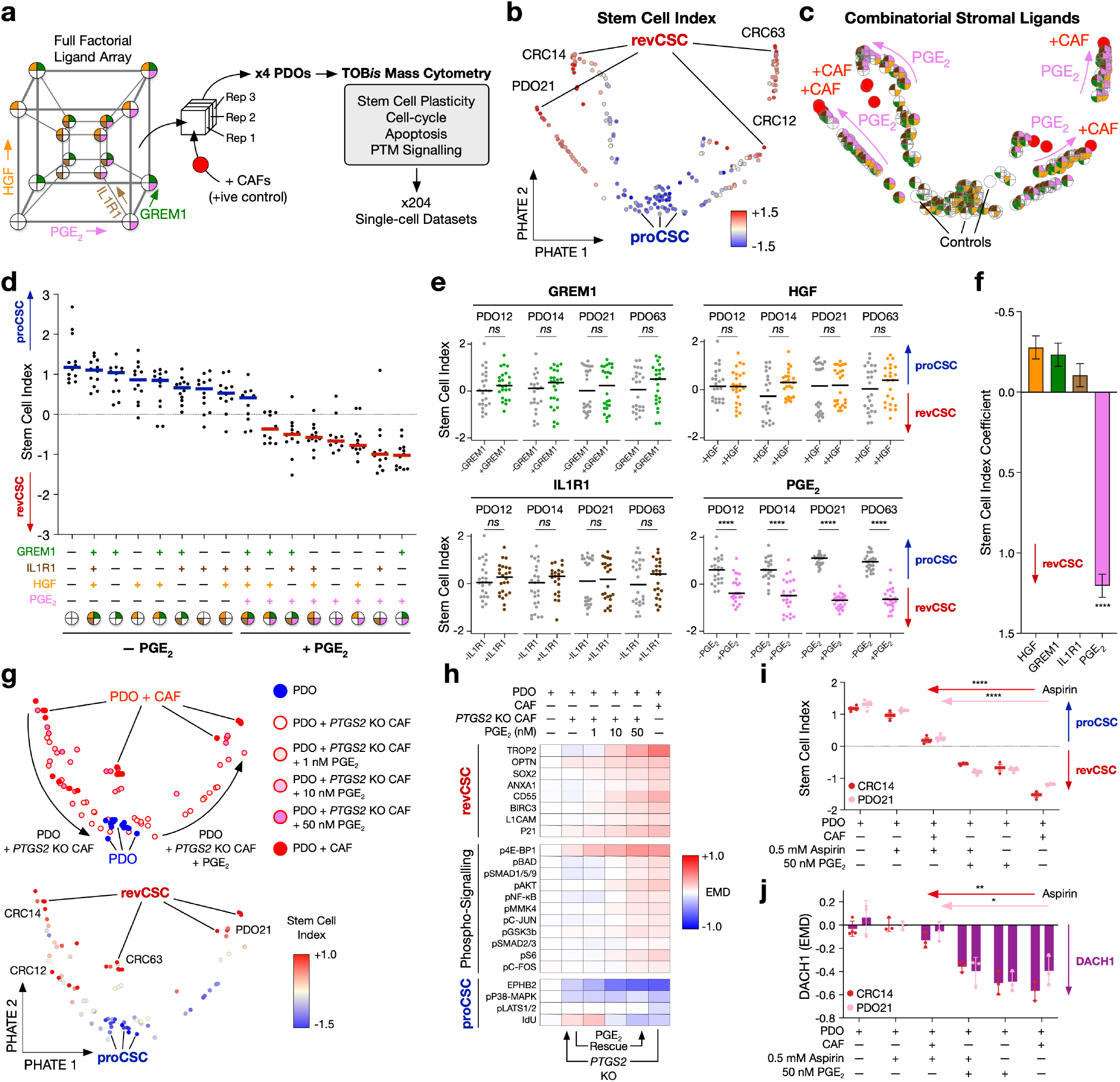
CAF-derived PGE_2_ is a Dominant Regulator of CRC Epithelial Plasticity. **a)** Experimental overview testing all ligand combinations identified by secretome CRISPR screening (full 2^4^ fixed-level factorial array) in triplicate across x4 PDOs. **b)** Paired TreEMD-PHATE coloured by stem cell index, or **c)** ligand treatment. 1 dot = 1 condition. **d)** Stem cell index of each ligand treatment across all PDOs. **e)** Stem cell index +/-each ligand per PDO. **f)** Linear mixed model effect coefficient of each CAF ligand on epithelial stem cell index. **g)** *PTGS2* KO CAFs co-cultured with x4 PDOs +/-1, 10, 50 nM PGE_2_ rescue. **h)** EMD heatmap of epithelial cell-fate and signalling markers following co-culture with *PTGS2* KO CAFs +/-1, 10, 50 nM PGE_2_ rescue. **i)** Stem cell index and **j)** DACH1 expression of PDOs co-cultured with CAFs +/-0.5 mM Aspirin +/-50 nM PGE_2_. Unpaired t-test, **** =p<0.0001, *** =p<0.001, ** =p<0.01, * =p<0.05, *ns* = not significant.

Single-cell treatment effects of each ligand combina-tion on each PDO were computed with paired TreEMD optimal transport [14] and embedded as a multi-PDO ligand-driven phenoscape. While each PDO displayed a distinct response to perturbation, all CRC cells formed a clear proCSC to revCSC transdetermination axis (Figure 5b) driven by all ligand combinations. Notably, we found that while IL1R1, GREM1, and HGF had a modest effect on PDO cell-fate, PGE_2_ treated epithelia phenocopied PDOs directly co-cultured with CAFs (Figure 5c, Figure S4a). When comparing the effect of all ligands across all possible combinations, we found that PGE_2_ consistently and significantly induced a proCSC to revCSC transition in all PDOs (Figure 5d, e). Linear mixed modelling analysis of all ligands across the full factorial array confirmed that PGE_2_ is a master effector of epithelial revCSC (Figure 5f). Collectively, these results strongly suggested that PGE_2_ is a dominant regulator of epithelial CRC plasticity.

To test if PGE_2_ is necessary and sufficient for CAF-induced proCSC to revCSC plasticity, we next CRISPR-engineered stable *PTGS2* knockout CAFs (Figure S4b, c), co-cultured them with x4 CRC PDOs +/-titrations of PGE_2_ and measured the single-cell PTM signalling and cell-fate responses by TOB*is* MC. This analysis revealed that *PTGS2* KO CAFs cannot alter the stem cell index of CRC cells, but proCSC to revCSC plasticity can be rescued by exogenous PGE_2_ (Figure 5g, h). Next, we treated CRC CAF-PDO co-cultures +/-aspirin (COX1/2 inhibitor) +/-PGE_2_ and measured epithelial single-cell signalling responses. Aspirin significantly reduced CAF-induced proCSC to revCSC plasticity and this effect could be rescued by exogenous PGE_2_ (Figure 5i). Finally, we found that both CAFs and exogenous PGE_2_ directly reduce epithelial DACH1 protein levels, whereas inhibiting CAF COX1/2 via aspirin can restore DACH1, even in the direct presence of *PTGS2*^*+*^ CAFs (Figure 5j).

To understand the dynamics of epithelial prostaglandin signalling, we performed a single-cell time-course phospho-protein, cell-cycle, and cell-fate analysis of CRC PDOs treated with PGE_2_. Time-course analysis revealed that PGE_2_ signals through epithelial pS6 [S240/S244], pP38 [T180/Y182], pNDRG1 [T346], p4E-BP1 [T37/T46], pC-JUN [S73], pMKK4 [S257], and pBAD [S112] in the first 6 hours (Figure S5). This initial signalling flux results in an early mitotic surge, that later transitions into a cell-fate switch with loss of DACH1, cell-cycle exit, and a plasticity transition from proCSC to revCSC. Interestingly, we observed that in the absence of additional PGE_2_ stimulation, CRC epithelia revert back to a more proCSC-like state – suggesting PGE_2_ acts as an acute, local, and reversible plasticity regulator in CRC. Collectively, these results confirm that CAF *PTGS2* is both a genetic and pharmacological dependency of CRC epithelial cell-fate and reveal that PGE_2_ is an acute master regulator of stromal-induced plasticity in CRC.

### *PTGS2*^*+*^ CAFs and Epithelial *DACH1* Map a Spatially Resolved Epithelial Plasticity Gradient in Human CRC

High-throughput single-cell perturbation analysis of PDO-CAF co-cultures revealed that *PTGS2*^*+*^ CAFs transdeterminate *DACH1*^*+*^ epithelia towards a revCSC fate. To explore whether this process is maintained in human tumours *in vivo*, we first analysed bulk RNA-seq data from human CRC samples. In agreement with *in vitro* organoid data, we found that *DACH1* is expressed in epithelia and *PTGS2* is expressed in stromal CAFs *in vivo* (Figure S6a,b). Concordantly, highly epithelial CMS2 tumours are *DACH1*^High^ whereas stromal CMS1/4 tumours are PTGS2^High^ (Figure S6c,d). Consistent with PGE_2_ suppressing *DACH1*, we find that *PTGS2* and *DACH1* anti-correlate in CRC tumours (Figure S6e) and *PTGS2*^*High*^/*DACH1*^*Low*^ tumours express more revCSC genes than *PTGS2*^*Low*^/*DACH1* ^*High*^ CRC tumours (Figure S6f). *PTGS2*^*High*^ tumours are more likely to be *BRAF*^*Mut*^, MSI, and CIN^Low^ whereas *DACH1*^*High*^ tumours are more commonly *KRAS*^*Mut*^, MSS, and CIN^High^ (Figure S6g). Collectively, these dichotomous results suggest that CAF *PTGS2* and epithelial *DACH1* anti-correlate in human CRC.

To explore the topographical relationships between CAF *PTGS2*, epithelial *DACH1*, and CRC stem cell plasticity we next analysed 5 primary CRC tumours with Xenium 5K spatial transcriptomics. In agreement with our *in vitro* perturbation studies, Topographical Correlation Map (TCM) analysis [22] revealed that *PTGS2*^*+*^ CAFs form a spatially resolved *DACH1* exclusion zone in CRC tumours *in vivo* (Figure 6a, b) (Figure S7a,b). Specifically, epithelia close to *PTGS2*^*+*^ CAFs are consistently *DACH1*^*–*^ whereas epithelia further away from *PTGS2*^*+*^ CAFs are *DACH1*^*+*^. We find that *PTGS2*^*+*^ CAFs exclude epithelial *DACH1* for approximately 200-300 µm (Figure 6c, d) (Figure S7c), consistent with the role of PGE_2_ as an acute short-range signalling molecule during wound healing [23].

**Figure 6.**
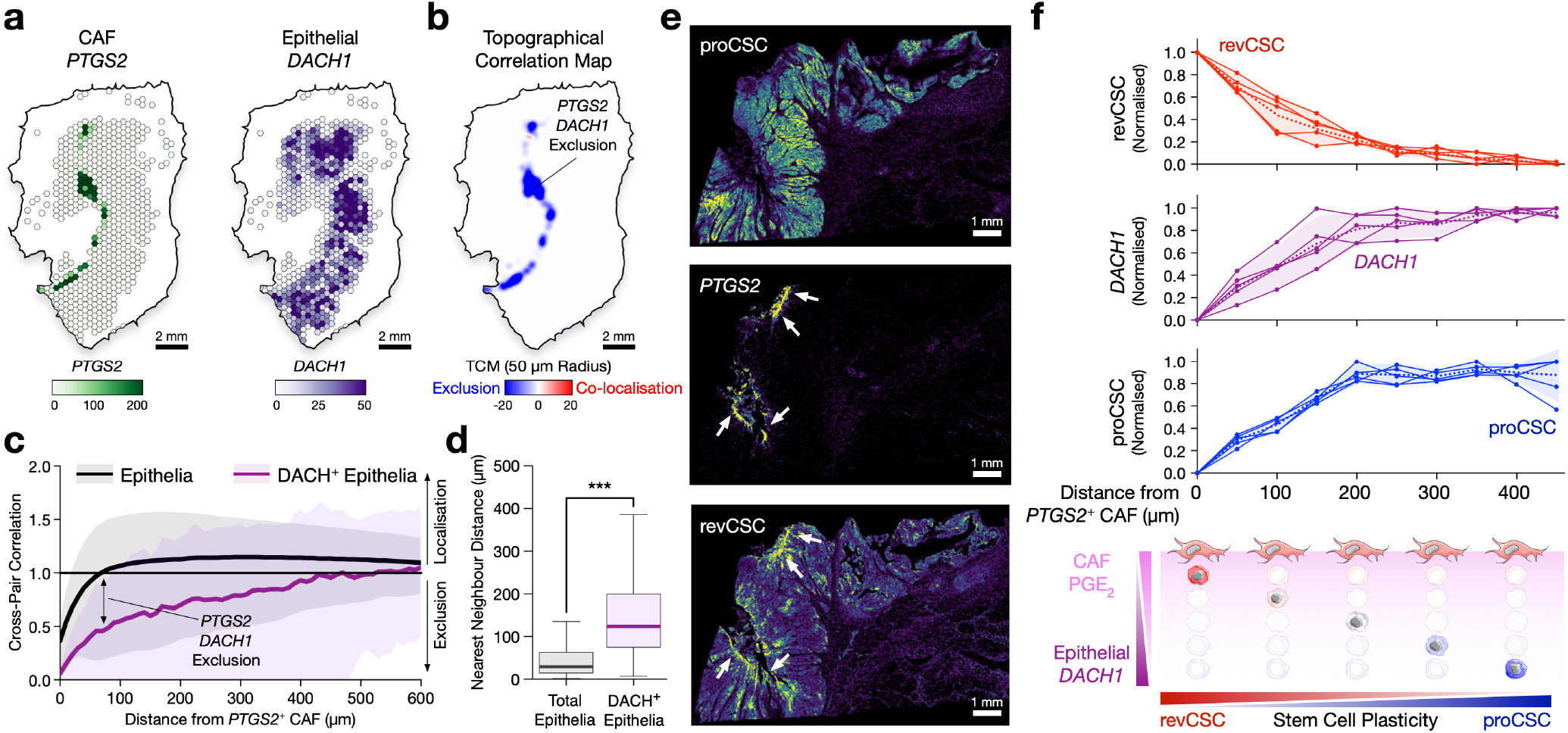
*PTGS2*^*+*^ CAFs and *DACH1*^*+*^ Epithelia Map a Spatially Resolved revCSC-to-proCSC Plasticity Gradient in Human CRC. **a)** CAF *PTGS2* and epithelial *DACH1* expression of a human primary CRC tumour analysed by Xenium 5K (hexgrid side length 200 µm). **b)** Topographical Correlation Map of *PTGS2*:*DACH1* co-localisation and exclusion (50 µm radius). **c)** Cross-pair correlation function analysis of all CAFs and all epithelia (black) versus *PTGS2*^*+*^ CAFs and *DACH1*^*+*^ epithelia (purple) from x5 CRC primary tumours. Mean, 95% CI. **d)** Nearest neighbour distance between all CAFs and all epithelia (black) versus *PTGS2*^*+*^ CAFs and *DACH1*^*+*^ epithelia (purple) from x5 CRC primary tumours. Mann-Whitney U test, *** = *p*<0.001. **e)** Transcript densities of *PTGS2* and proCSC and revCSC gene signatures in a human CRC tumour. **f)** *DACH1*, proCSC and revCSC gene signatures relative to *PTGS2*^*+*^ CAFs from 5 CRC tumours. Dashed line = mean, fill = 95% CI.

To understand whether the *PTGS2*-*DACH1* gradient correlates with CRC stem cell-fate, we next studied proCSC and revCSC gene expression signatures relative to *PTGS2*^*+*^ CAFs (Figure 6e) (Figure S7d). Across all tumours, we observed a highly reproducible, spatially resolved revCSC to proCSC gradient relative to *PTGS2*^*+*^ CAFs. Specifically, epithelia <200 µm from *PTGS2*^*+*^ CAFs are enriched for revCSC, whereas epithelia >200 µm from *PTGS2*^*+*^ CAFs are proCSC dominant (Figure 6e, f). Moreover, the revCSC to proCSC axis spatially aligns with *DACH1* expression in all CRC tumours. In summary, by combining systematic single-cell perturbation analysis, intercellular CRISPR screening, and spatial transcriptomics, these results collectively suggest that stromal prostaglandin is a master regulator of epithelial stem cell plasticity in human CRC.

## Discussion

Stromal CRC tumours have poor response to therapy, increased metastasis, and poor prognosis when compared to highly epithelial tumours [12, 13]. While multiple stromal-epithelial interactions have been identified [2, 10], the master determinants of intercellular signalling in CRC tumours have remained unclear. Here we applied high-throughput single-cell perturbation analysis of human CRC PDO-CAF co-cultures, to explore the molecular regulators of epithelial cell-fate regulation by CAFs. We find that CRC cells occupy a fundamental proCSC to revCSC plasticity axis that correlates with standard-of-care chemotherapy responses. proCSC are exquisitely chemosensitive, whereas revCSC are chemorefractory. We find that not all CRC cells respond to stromal cues from CAFs, but epithelia with high-plasticity potential express *Dachshund homolog 1* (*DACH1*). DACH1 is a transcriptional co-repressor that can interact with AP-1 [24], NF-κB [25, 26], YB-1 [27], SMAD4 [28], and SNAI1 [29] transcription factors, and has been shown to regulate β-catenin [30], BMP [31], and TGF-β signalling [32] across a range of cell-types. In *Drosophila*, null mutations in *dachshund* result in flies with no eyes and shortened legs [33] – resembling Dachshund ‘sausage dogs’ – underscoring its important role in cell-fate determination. In mammals, DACH1 has been shown to play a role in epithelial response to inflammation in the lung [34, 35] and regulate cell-fate in breast cancer [27]. However, the role of DACH1 in CRC has not been widely explored. Previous research has shown that *DACH1* is overexpressed in CRC, can localise to colonic crypt base cells, and predicts poor prognosis [31, 36]. However, DACH1 has also been described as a suppressor of tumour growth and metastasis in CRC [27, 37, 38]. These counter-intuitive observations suggest that DACH1 may not represent a classical tumour promoter or suppressor – but instead DACH1 plays a more dynamic role in CRC biology. We find that *DACH1* is not associated with a single attractor state in CRC, but is instead expressed in subsets of proCSC, revCSC, squamous, and neuroendocrine cells. This broad expression may explain why the role of *DACH1* has previously been paradoxical. However, we find that instead of being associated with a single cell-type, *DACH1* is a single-gene predictor of whether CRC cells can communicate with CAFs. *DACH1* does not define a single cell-fate, but rather marks epithelial competence for stromal reprogramming.

Dozens of intercellular signalling molecules have been implicated in CRC tumour biology [2, 10]. However, the relative functional contribution of individual ligands has not been defined. Through single-cell perturbation analysis of the CAF secretome, we demonstrate that stromal regulation of *DACH1*^*+*^ epithelia can occur via *HGF, IL1R1, GREM1*, and *PTGS2* – but mesenchymal *PTGS2* is a dominant regulator of CRC epithelial fate relative to other ligands. Over 30-years ago Eberhart *et al*. identified that *PTGS2* (COX2) is overexpressed in human CRC tumours [39] and subsequent research has revealed that COX2, and its soluble product PGE_2_, play an important role in CRC – particularly in tumour initiation. For example, stromal PGE_2_ regulates CRC tumour initiation in mice [40], *PTGS2*^*+*^ CAFs are found in early-stage CRC [41], and pharmacological COX2 inhibition can block CRC tumour formation [42]. Both aspirin (COX1/2 inhibitor) [43] and celecoxib (COX2 inhibitor) [44] can reduce CRC incidence in humans. PGE_2_ has also been shown to regulate epithelial wound healing [45], promote CRC stem cell expansion in mice [46], and regulate proliferation via PI3K and β-catenin signalling [47]. These studies all demonstrate that COX2 and PGE_2_ are important for tumour initiation and cancer cell proliferation.

However, these studies do not explain why COX2 overexpression is associated with worse survival in established human CRC [48]. Here, through single-cell perturbation analysis of the CAF secretome, we demonstrate that PGE_2_ is a dominant stromal regulator of CRC epithelial plasticity. More than any other native ligand from the CAF secretome, stromal PGE_2_ rapidly induces a revCSC phenotype in CRC epithelia, which is strongly associated with poor chemotherapy response, metastasis, and disease relapse [10]. We find that *PTGS2*^*+*^ CAFs form a spatially-resolved stem cell transdetermination gradient in human tumours, providing localised non-genetic access to poor-prognosis revCSC cell-fates. Rather than viewing revCSC as a stable intrinsic cell-fate, our data suggest that revCSC identity is locally regulated by *PTGS2*^*+*^ stromal niches. *PTGS2*^*+*^ CAFs therefore permit acute non-genetic access to a pro-metastatic, chemoresistant cell-fate with access to further non-canonical differentiation [9]. Crucially, we show that genetic and pharmacological inhibition of stromal PGE_2_ blocks epithelial plasticity. These results may explain why low-dose aspirin significantly lowers the incidence of CRC recurrence among patients with PIK3CA hotspot mutations [49], CRC patients who take aspirin have fewer metastases [50], and adjuvant celecoxib can improve survival in ctDNA^+^ patients irrespective of PIK3CA status [51]. Our data suggest that COX inhibitors do not act on epithelial cells directly, but instead reduce stromal-induced access to revCSC – which is required for metastasis and chemoresistance. Under this model anti-COX therapies can be considered to be intercellular anti-plasticity drugs.

In summary, previous studies have established that stromal CRC tumours have a worse outcome [12, 13]. In parallel, COX2/PGE_2_ has been shown to promote CRC initiation [40] and proliferation [46] in mice and anti-COX therapies have shown anti-CRC efficacy in humans [49, 50, 51]. Here, we mechanistically unite these observations by demonstrating that stromal PGE_2_ is a dominant spatial regulator of epithelial plasticity in human CRC. Stromal PGE_2_ drives an acute, transient, and reversible transition to a chemoresistant pro-metastatic cell-fate that is marked by decreased epithelial *DACH1* expression. These results provide a mechanistic rationale for anti-PGE_2_ strategies in both the prevention and treatment of CRC.

## Methods

### Protocols

#### CRC PDO and CAF Culture

CRC PDOs were obtained from the Human Cancer Models Initiative (Sanger Institute, Cambridge, UK) [52] and expanded in 3x 25 µL droplets of Growth Factor Reduced Matrigel (Corning 354230) per well of a 12-well plate (Helena Biosciences 92412T). Each well was supplemented with 1 mL of PDO expansion media comprising Advanced DMEM F/12 (Thermo 12634010) containing 2 mM GlutaMAX (Thermo 35050061), 1 mM N-acetyl-L-cysteine (Sigma A9165), 10 mM HEPES (Sigma H3375), 500 nM A83-01 (Generon 04–0014), 10 µM SB202190 (Avantor CAYM10010399-10), and 1X B-27 Supplement (Thermo 17504044), 1X N-2 Supplement (Thermo 17502048), 50 ng mL^−1^ EGF (Thermo PMG8041), 10 nM Gastrin I (Sigma SCP0152), 10 mM Nicotinamide (Sigma N0636), 1X HyClone Penicillin-Streptomycin Solution (Fisher SV30010), and conditioned media produced as described in [53] at 5% CO_2_, 37°C. L-cells for conditioned media production were obtained from Shintaro Sato (Research Institute of Microbial Diseases, Osaka University, Osaka, Japan) [54]. PDOs were dissociated into single cells with 1X TrypLE Express Enzyme (Gibco 12604013) (incubated at 37°C for 20 min) and passaged every 5-10 days. For the first 24 hours after dissociation, media was also supplemented with 10 µM Rho-associated protein kinase inhibitor (ROCKi) (Y-27632, Sigma Y0503). CRC CAFs were a kind gift from Prof. Olivier De Wever (University of Gent) [55]. All CAFs were cultured in DMEM (Thermo 11965092) enriched with 10% FBS (Gibco 10082147), and 1X HyClone Penicillin-Streptomycin Solution (Fisher SV30010) at 5% CO_2_, 37°C.

For experimental analysis, PDO were dissociated into single cells on day 1 and expanded in 5% Matrigel solution of Advanced DMEM F/12 (Thermo 12634010) containing 2 mM GlutaMAX (Thermo 35050061), 1 mM N-acetyl-L-cysteine (Sigma A9165), 10 mM HEPES (Sigma H3375), 1X B-27 Supplement (Thermo 17504044), 1X N-2 Supplement (Thermo 17502048), 50 ng mL^−1^ EGF (Thermo PMG8041), 10 nM Gastrin I (Sigma SCP0152), 10 mM Nicotinamide (Sigma N0636), 500 nM A83-01 (Generon 04–0014), 10 µM SB202190 (Avantor CAYM10010399-10), and 1X HyClone Penicillin-Streptomycin Solution (Fisher SV30010) in ULA T75 flasks (Corning BC371) at 5% CO_2_, 37°C for 4 days. On day 5, PDOs were pelleted and transferred into base media (Advanced DMEM F/12 containing only 2 mM GlutaMAX, 1 mM N-acetyl-L-cysteine, 10 mM HEPES, 1X B-27 Supplement, 1X N-2 Supplement, 10 mM Nicotinamide, and 1X HyClone-Penicillin Streptomycin Solution). In parallel, CAFs were starved in 2% FBS DMEM with 1X Hyclone-Penicillin Streptomycin Solution. Unless otherwise described culture conditions are as follows. PDOs and CAFs were seeded on day 6 in 96-well plates (Helena Biosciences 92696T) in 50 µL Matrigel with 300 µL base media. PDOs are seeded at a density of ~1.5 x 10^3^ organoids per well and CAFs at 1.0 x 10^5^ cells per well. Cultures were maintained for 72 hours with media refreshed every 24 hours.

#### scRNA-seq of CRC PDOs

scRNA-seq of CRC PDOs was performed using the SPLiT-seq method described in Ramos Zapatero *et al*. [14]. In brief, PDOs were cultured in triplicate as described above for 72 hours with fresh reduced media replaced every 24 hours. PDOs were harvested and dissociated into single cells using TrypLE (Thermo 12604013) incubated for 10 minutes at 37°C on a heated orbital shaker at 300 rpm. Cells were filtered through a 35 µM filter and re-suspended in 1 mL of PBS supplemented with 1.25 µL Protectorase RNase inhibitor (Merck 3335402001) and 2.5 µL Superase RNase inhibitor (Thermo AM2694). Cells were fixed, permeabilised, counted and 5% (v/v) DMSO was added before aliquoting and freezing in a Mr Frosty at −80°C. One complete SPLiT-seq experiment was performed per two PDOs studied (five independent SPLiT-seq runs total). Split-pool barcoding combinatorial indexing was performed as per the SPLiT-seq protocol [56] with minor modifications. Briefly, cells were thawed and reverse transcribed in the barcode-RT1 plate, followed by two rounds of pooling and ligation in the ligation-L2 and ligation-L3 plates. Cells were then counted and aliquoted into sublibraries of approximately 12,500 cells, lysed and frozen at −80°C until library preparation.

Following split-pool barcoding, 4 sub-libraries were carried forwards for cDNA isolation, amplification and library generation as described in Ramos Zapatero *et al*. [14]. Libraries were then pooled together equally, loaded onto an Illumina Novaseq (200 cycle NovaSeq 6000 S2 Reagent Kit v1.5) and sequenced within the following format: R1:74-i7:06-i5:00-R2:86 to yield a paired end read structure comprising cDNA transcriptomic information in read 1 and barcoding information in read 2. The data were demultiplexed using the sublibrary i7 indexes and reads aligned to the GRCh38 reference genome using the zUMIs package (v2.9.7) [57] with STAR (v2.7.3a), filtering on a whitelist of permitted cell barcodes and merging cells that shared PolyA and Random Hexamer RT1 barcodes from the same RT well plate position with identical L2 and L3 barcodes, including reads originating from exons and introns. Cell barcodes were collapsed based on a 2 hamming distance of close cell barcodes and UMIs on a 1 hamming distance of UMI sequence. A cell x gene digital gene expression matrix (DGE) was generated for each library.

#### scRNA-seq Data Preprocessing and Analysis

For all sequencing runs, the DGE of each sublibrary was processed using the splitRtools package (https://github.com/TAPE-Lab/splitRtools) to annotate cell barcode well locations and sample identities, and perform initial QC. Downstream analysis was performed in Scanpy [58]. Sublibrary DGEs were merged and low quality cells excluded using the following parameters; >32,500 unique molecular identifiers (UMIs) and <1000 UMIs, >7000 genes and <400 genes, >20% mitochondrial transcripts, >0.4 UMI/read ratio and genes detected in fewer than 25 cells. Neotypic doublets were identified and removed using Scrublet [59] with an expected doublet rate of 3% based on previous published data. Cells were then normalised using count based normalisation with a scaling factor of 10,000 excluding highly expressed genes comprising >5% of total counts per cell. The normalised data were then natural log transformed for downstream analysis. Data were scaled and PCA performed over 5000 variable genes, a neighbourhood graph was constructed based on the first 50 PCs as input and Leiden clustering performed to generate clusters. Once all scRNA-seq datasets had been filtered as above, the final DGEs were merged, ultimately yielding a dataset of 160,898 high-quality single cells and 39,835 genes.

##### Integration

Data integration was performed using Scanpy’s harmony_integrate implementation with default parameters [60] to integrate single-cell data from multiple experiments. Integration was computed using all cells, PDO epithelial cells from monoculture or mono- and co-culture conditions, CAF cells from monoculture or mono- and co-culture conditions.

##### Dimensionality Reduction

For dimensionality reduction (DR), the first 50 PCs were computed from the integrated objects. For the PHATE embeddings, 50, 75, or 100 PCs were used with default parameters. PHATE was chosen as the standard DR method for data visualisation throughout this study due to its capacity to capture global dataset architecture in biological systems with important developmental trajectories [61].

##### Gene Signature and Cell Cycle Scoring

All gene expression signature scores were computed using the score_-genes function in Scanpy using default settings on Log-Normalised data. Scores were computed using a defined set of genes and comparing their average relative expression against a reference set of genes, as previously described in Satija *et al*. [62]. proCSC and revCSC signatures were from Opzoomer *et al*. [63]. Squamous and Neuroendocrine signatures were derived from Moorman *et al*., [9] and Goblet signatures were from Malla *et al*., [6]. Cell cycle scores were computed using the score_-genes_cell_cycle function in Scanpy using a curated list of cell cycle genes from Tirosh *et al*. [64].

##### MELD Likelihood Analysis

MELD was used to establish cells (PDOs or CAFs) within each culture condition that had been perturbed by the CAF co-culture condition. MELD is a manifold-geometry based method used to quantify the effect of an experimental perturbation (e.g., + CAFs) by estimating the relative likelihood of observing cells in each experimental condition over a graph learned from all cells in a sample [16]. To preserve enrichment information across replicates, a density estimate was generated per replicate and then L^1^ normalisation was applied to these densities within each replicate to normalise the values to a sum of 1 across samples within each replicate. Average likelihoods of PDO-CAF co-culture samples were utilised as the measure of perturbation. To calculate a single score per PDO, the MELD likelihood delta was calculated by subtracting the mean likelihood score of monoculture replicates from the mean likelihood score of co-culture replicates on a per PDO basis.

##### Differential Gene Expression

Differentially expressed genes (DEGs) between conditions were identified using the Wilcoxon rank-sum test with P-values adjusted for multiplicity of tests using Benjamini-Hochberg adjustment implemented by Scanpy’s rank_gene_groups function. Volcano plots were plotted for the top DEGs between conditions selected with adjusted P-value <0.01 and an absolute value Log Fold Change >1.

##### Stem Cell Index Calculation

Using the merged dataset PDO specific proCSC and revCSC signature scores were computed for each PDO epithelial cell using literature derived proCSC and revCSC signatures. Following gene scoring, single-cell gene signature scores were Z-score normalised, and the stem cell index calculated by subtracting the proCSC score from the revCSC score to determine the relative distribution of different stem cell signatures within each PDO, as described in Vasquez *et al*. [5].

##### Pluripotency Measure

The pluripotency values for PDO epithelial cells were calculated using the R package Cyto-TRACE 2 [65], which leverages the concept that differentiated cells exhibit lower transcriptional gene expression and diversity compared to cells with high potency.

##### Transcription Factor Activity Inference

Transcription factor activity analysis was performed using the decou-pleR Python package [15] using the Omnipath CollecTRI knowledge graph network [66], comprising an extensive resource of curated transcription factors (TFs) and their associated gene targets. Briefly, a linear model is fitted to estimate the observed gene expression based on a given TF’s TF-gene interaction weights. The t-value of the slope is then taken as the TF activity score. TF activities were compared across conditions using the decoupler rank_sources_groups function with the Wilcoxon rank-sum test to compute differences between groups.

##### Valley Ridge Landscapes

VR landscapes were computed and visualised as described in Qin *et al*., [3]. All gene signatures are available in Supplementary File ‘CRC_-scRNA-seq_Modules_All.csv’.

#### Thiol-reactive Organoid Barcoding *in situ* (TOB*is*) Mass Cytometry

PDO monocultures, CAF monocultures and PDO-CAF co-cultures were analysed using the TOB*is* mass cytometry protocol described in Sufi & Qin *et al* [20]. In brief, PDO-CAFs cultures were incubated with 25 µM (5-Iodo-20-deoxyuridine) (^127^IdU) (Standard BioTools 201127) at 37°C for 30 min, and 5 min before the end of this incubation, 1X Protease Inhibitor Cocktail (Sigma, P8340) and 1X PhosSTOP (Sigma 4906845001) were added into the media. Each well was then fixed in 4% PFA/PBS (Thermo J19943K2) for 1 hour at 37°C. PDO-CAFs were washed with PBS and *in situ* with 126-plex (9-*choose*-4) TOB*is* overnight at 4°C. Unbound barcodes were quenched in 2 mM GSH and all PDO-CAFs were pooled. PDO-CAFs were dissociated into single cells using 1.0 mg mL^−1^ Dispase II (Thermo 17105041), 0.2 mg mL^−1^ Collagenase IV (Thermo 17104019), and 0.2 mg mL^−1^ DNase I (Sigma DN25) in M-Tubes (Miltenyi 130-093-236) via gentleMACS Octo Dissociator with Heaters (Miltenyi 130-096-427, SCR_020271). Single PDOs and CAFs were washed in cell staining buffer (CSB) (Standard BioTools 201068) and stained with extracellular rare-earth metal conjugated antibodies (Table S1) for 30 min at room temperature. PDO-CAFs were then permeabilised in 0.1% (v/v) Triton X-100/PBS (Sigma T8787) and stained with intracellular rare-earth metal conjugated antibodies overnight at 4°C. PDO-CAFs were then washed in CSB followed by distilled H_2_O and incubated for 10 min in 10 mg mL^−1^ Ruthenium 0.1 M NaHCO_3_ solution (Sigma 96631) for cell volume normalisation [67]. After CSB wash, PDO-CAF were then incubated in 125 nM^191^ Ir/^193^ Ir DNA intercalator (Standard BioTools 201192A) for 1 hour at room temperature. PDO-CAFs were washed, resuspended in cell acquisition solution plus (CAS+) (Standard BioTools 201244), and analysed using a CyTOF XT (Standard BioTools, SCR_02634) at 200–400 events s^−1^.

#### CRC PDO Chemosensitivity Assay

PDOs were cultured as described above and seeded into clear-bottomed 96-well plates (Greiner, 655098) in technical triplicates. Cultures treated with 10 doses of 5-FU, oxaliplatin and SN-38, as well as a lethal control of 50% DMSO and a vehicle control. 48 hours after treatment, all media was removed and 50*µ*L CellTiter-Glo 3D (Promega, G9681) was used to assess viability according to the manufacturer’s protocol. Luminescence readings were obtained using the Varioskan Lux plate reader (Thermofisher).

#### Mass Cytometry Data Processing

Multiplexed FCS files were debarcoded into discrete experimental conditions using the Zunder Lab Single Cell Debarcoder (https://github.com/zunderlab/single-cell-debarcoder) [68]. Debarcoded FCS files were uploaded to Cytobank and gated for Gaussian parameters and DNA content (^191^Ir/^193^Ir). Epithelial PDO cells were gated as Pan-CK^+^ EpCAM^+^ and CAFs gated as Vimentin^+^ FAP^+^. If required batch normalisation was conducted using cytoBatchNorm (https://github.com/i-cyto/cytoBatchNorm) and exported PDO and CAF cells were cell volume normalised [67] and arcsinh transformed prior to downstream analysis.

#### Geodesic Sinkhorn Optimal Transport Analysis

Geodesic Sinkhorn is a fast, memory-efficient, geometry-aware method for solving the entropy-regularised optimal transport (OT) problem between distributions *µ* and *v* that lie on or near a manifold [21],

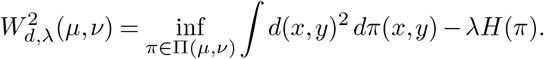

The geodesic distance on the manifold is the natural ground cost, but computing it directly is generally infeasible. Varadhan’s formula resolves this by expressing geodesic distance as a limit of heat diffusion at small timescales,

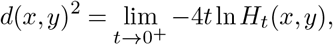

where *H*_*t*_ = *e*^*−tL*^ is the heat kernel. This motivates the heat-geodesic ground distance 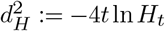. Setting *λ* = 4*t*, the entropy-regularised problem becomes a KL projection onto the heat kernel:

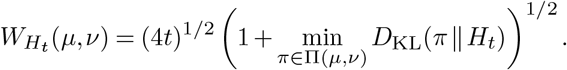

To compute this, an undirected graph *G* = (*V, E*) is built over the data points from the union of distributions and its sparse Laplacian *L* = *D − A* is formed. The heat operator *H*_*t*_ = *e*^*™tL*^ is approximated by a Chebyshev polynomial *p*_*K*_(*L, t*) of degree *K*, with approximation error decaying exponentially in *K*. Sinkhorn iterations then update scaling vectors *v, w* via

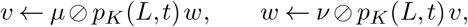

requiring only sparse matrix-vector products. At convergence, the optimal coupling 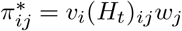 and the marginal constraints 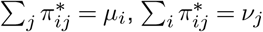 collapse the KL term to

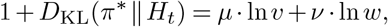

yielding the entropy-regularised Wasserstein distance from the converged scaling vectors. For single-cell data application, these distances are computed pairwise between experimental conditions to populate a distance matrix.

#### Earth Mover’s Distance (EMD) Optimal Transport Analysis

Earth mover’s distance (EMD) scores were calculated using the Python package *scprep* (https://scprep.readthedocs.io/en/stable/index.html). Standard EMD scores were calculated between probability distributions (markers) across experimental conditions using defined controls as references. Signs were applied to EMD scores to denote an increase or decrease in expression.

#### Determining Stem Cell Index (SCI) from TOB*is* Mass Cytometry Data

Markers representing the proCSC state (EPHB2, pLATS1, TOP2A, pP38-MAPK, IdU, CDK1 & pRB) and the revCSC state (TROP2, CD55, ANXA1, BIRC3, CK18, OPTN & SOX2) were first z-scaled to ensure each marker contributed to the score equally before a mean of each group was calculated and the revCSC score is subtracted from the proCSC score. For TOB*is* MC data, SCI was oriented such that positive values represent proCSC enrichment and negative values represent revCSC enrichment, as indicated in each figure. When multiple PDO lines were used the SCI score per PDO was standardised to enable comparisons between PDOs. To statistically evaluate ligand dominance across the 2^4^ factorial array, per-PDO standardised SCI scores were modelled using a linear mixed-effects model with ligand presence/absence as fixed effects and PDO line as a random intercept. Fixed-effect coefficients and p-values were estimated using Wald tests (Python statsmodels MixedLM).

#### PHATE

Potential of heat diffusion for affinity-based transition embedding (PHATE) is a non-linear dimensionality reduction method that aims to preserve the local and global structure of datasets [61]. A variety of data matrices can drive PHATE embeddings, including EMD scores and ΔDREMI scores, whereby the resulting embedding captures maximal variance across all the dataset dimensions. Conclusions can be made surrounding the relative distance of points within a PHATE embedding, which are linked to underlying similarities or differences in the chosen markers of a mass cytometry panel (as captured by EMD calculation).

#### Computational Analysis of Human CRC Cohorts

Microarray gene expression data from the human CRC cohort were accessed from Gene Expression Omnibus (GSE39582) [69]. Probes were collapsed to gene level using the maximum mean method. Analysis was restricted to untreated patients (no adjuvant chemotherapy; n = 316). proCSC and revCSC gene module scores were computed using single-sample gene set enrichment analysis (ssGSEA) implemented in GSVA (v2.4.4), with a minimum gene set size of 3 and score normalisation enabled. A stem cell index was defined as revCSC minus proCSC score. Tumours were stratified into *PTGS2*-high/*DACH1*-low and *PTGS2*-low/*DACH1*-high groups using median expression as the threshold for each gene independently. Group comparisons were performed using the Wilcoxon rank-sum test. All analyses were performed in R (v4.5.2) using ggplot2 and ggpubr. UMI count data for the SMC CRC cohort (GSE132465; [18] n = 23 patients) were obtained from the Gene Expression Omnibus and processed using Seurat (v4.4.0). Genes detected in fewer than 3 cells were excluded. Low-quality cells were further removed based on the following thresholds: fewer than 200 or more than 6,000 detected features, or mitochondrial gene content exceeding 25%. Analysis was restricted to tumour epithelial cells (n = 17,469). Data were normalised using log-normalisation, and the top 2,000 highly variable genes were identified using the variance-stabilising transformation method. Data were scaled and principal component analysis (PCA) was performed on variable features. Patient-specific batch effects were corrected using Harmony (v2.0.3) applied to the first 30 principal components. Diffusion map analysis was performed on the first 10 Harmony-corrected principal components using the destiny R package (v3.24.0). The first diffusion component (DC1) was used as a trajectory axis, along which cells were ordered into 100 equal bins. Mean *DACH1* expression per bin was smoothed using a rolling mean with a window of 5 bins. For visualisation, PHATE dimensionality reduction was applied to the first 10 Harmony-corrected principal components using phateR (v1.0.7, KNN = 15). Gene expression programme scores for proCSC, revCSC, Goblet, Neuroendocrine, and Squamous modules were calculated per cell using AddModuleScore() in Seurat. Each cell was assigned to the module with the highest score, provided that score exceeded the 67th percentile threshold for that module; cells not meeting this threshold for any module were labelled Unassigned.

The bulk RNAseq of human CRC cohort were accessed from Gene Expression Omnibus with accession numbers GSE39582. The probes-to-genes were collapsed using the collapseRows function in the WGCNA R package (v.1.70-3), in which the probe with the highest average value per gene was selected. Analysis was restricted to untreated patients (no adjuvant chemotherapy; n = 316). *DACH1* and *PTGS2* expression values were derived from microarray data (log2-transformed, probe-collapsed to gene level by maximum mean). Tumours were stratified into *PTGS2*-high/*DACH1*-low and *PTGS2*-low/*DACH1*-high groups using median expression as the threshold for each gene independently. proCSC and revCSC gene module scores were computed as previously described, and a stem cell index was defined as revCSC minus proCSC score. All analyses were performed in R (v4.5.2) using ggplot2 and ggpubr.

#### Paired TreEMD Optimal Transport Analysis

Paired TreEMD (Tree-Earth Mover’s Distance) is used to measure treatment-induced changes between single-cell distributions [14]. A hierarchical tree is built over the cell-feature space by recursive k-means clustering, each distribution is embedded as a vector of edge-weighted subtree masses, and matched-control embeddings are subtracted from each treated sample prior to taking the L1 norm yielding a control-corrected distance that captures the difference in treatment effect between conditions.

#### CAF-Conditioned Media Experiments

CRC CAF derived conditioned media was prepared by growing 25 x 10^6^ CAFs in 5% Matrigel/base media solution in ULA T75 (Corning BC371) for 72 hours. CAF-conditioned media was then filtered (Sartorius 10109180). PDO and CAFs were prepared for this study as described above. PDOs were treated with 0-5% CAF-conditioned media prepared in base media, the media was replaced every 24 hours for 72 hours. The cultures were processed via TOB*is* Mass Cytometry.

#### Generation of Cas9^+^ CRC CAFs

CRC CAFs were transduced with pLenti-Cas9-GFP vector (Addgene 86145) at a multiplicity of infection of 5 in 10% FBS DMEM supplemented with 8 µg/ml polybrene (Merck, TR-1003-G) and incubated overnight at 5% CO2, 37°C. CAFs were then maintained until sufficient number for cell sorting. Once enough cells acquired, CAFs were harvested and washed in FACS buffer (PBS supplemented with 1 mmol/L EDTA^+^(Sigma, 03690), 25 mmol/L HEPES, and 1% FBS) before sorting for GFP expression through a BD FACSAria III. Cas9 expression was then assessed by harvesting and fixing the CAFs in 4% PFA/PBS (Thermo J19943K2) on ice for 15 min before permeabilising the cells in 0.1% (v/v) Triton X-100/PBS (Sigma T8787). Cells were stained with rare-earth metal conjugated antibodies against Cas9, Vimentin and GFP (Table S1) and prepared for running on the CyTOF XT as described previously.

#### Intercellular CAF-secretome CRISPR Screening

CRISPR guide RNA (gRNA) library was prepared by resuspending the custom 0.1 nmol Edit-R pool crRNA library (Horizon) (see Supplementary File ‘Secretome_-CRISPR_Library.xls’) in 2.5 µM Edit-R tracrRNA (Horizon U-002005-20) in 10 mM Tris buffer (Horizon B-006000-100) and incubated for 30 min at room temperature. Edit-R crRNA controls, non-targeting control #2 (Horizon, U-007502-01-05), lethal control #1 (Horizon, U-006000-01-05) and B7-H3 (Horizon, CM-007813-02-0002), and Edit-R tracrRNA were reconstituted to 50 µM with 10 mM Tris and duplexed by combining at a 1:1 ratio and incubating at room temperature for 30 min before loading into library plate. The library was stored at −80°C.

On day 1 of the assay, duplexed gRNA were transferred into 96 well round bottomed assay plates (Thermo 163320) using an Echo 525 Acoustic Liquid Handler (Beckman Coulter 001-10080) to achieve a final assay concentration of 25 nM. Then, 20 µL of OptiMEM (Thermo 31985062) containing 0.4 µL Lullaby lipid transfection reagent (Oz Biosciences LL71000) was dispensed into each well and incubated for 20 min at room temperature. 80 µL Cas9 expressing CRC CAFs suspended in 2% FBS DMEM were then dispensed at a density of 4000 cells per well and the plates were maintained at 5% CO2, 37°C for 120 hours. On day 6, media was removed and CAFs detached using 1X TrypLE Express Enzyme. CAFs were pelleted by centrifugation at 200 G for 5 min and supernatant removed. CAF pellets were then resuspended in 50 µL Growth Factor Reduced Matrigel containing PDO (PDO cultured for experimentation as described previously) and 200 µL base media was added after Matrigel polymerisation, the media was refreshed every 24 hours. After 48 hours (day 8) of co-culture the PDO-CAF were processed via the TOB*is* Mass Cytometry protocol.

#### Evaluation of CRISPR Screen Knockout Efficiency by ELISA

Edit-R crRNA targeting *HGF* (Horizon CM-006650-01-0002), *CXCL8* (Horizon CM-004756-01-0002), *VEGFA* (Horizon CM-003550-02-0002) and *PTGS2* (Horizon CM-004557-02-0002) sequences: TAGTACATCTATTAGCACAT, CAGAGCTGCAGAAATCAGGA, GGAGGGCA-GAATCATCACGA & TGTACCCGGACAGGATTCTA and Edit-R tracrRNA was reconstituted to 50 µM with 10mM Tris and duplexed by combining at a 1:1 ratio and incubating at room temperature for 30 min. The protocol outlined above was followed and once the CAFs had been edited they were resuspended in Matrigel containing no PDO. Conditioned media from those cultures was harvested at 24 hours and 48 hours and assessed for protein expression via ELISA (Abcam ab275901, ab214030, ab222510, ab287802).

#### CRC PDO Isolation and Expansion

CRC patient tissue samples not required for diagnostic purposes were obtained from the UCL/UCLH Biobank for Studying Health and Disease following ethical approval (REC reference: 20/YH/0088). Tissue acquisition, storage, and use complied with institutional governance and, where applicable, the Human Tissue Act 2004. Written informed consent for tissue donation and research use was obtained from all participants before sample collection. PDOs were established at the Translational Technology Organoid Platform under the registered biobank project NC12.14. All surgical resections were transported in cold DPBS (Gibco) from UCLH, examined, and tumour tissue was sampled by a pathologist. Tumour samples (0.5 cm^3^ in size) were stored in Belzer UW Cold Storage Solution (Bridge to Life) supplemented with 0.1 mg/mL Primocin (InvivoGen) for either processing within 24 hours or longer-term storage in liquid nitrogen. Down-stream sample processing included tissue washing with DPBS and cutting into 2 mm^3^ fragments, which were subsequently enzymatically digested in 10 mL digestion buffer (1X Advanced DMEM base medium (Gibco), Collagenase II [5 mg/mL final] (Gibco), Deoxyribonuclease I [final 0.2 mg/mL] (Merck), Primocin [final 0.1 mg/mL] (InvivoGen), and Rho-associated protein kinase inhibitor Y-27632 [final 10 µM] (Sigma-Aldrich)). Samples were mechanically processed by placement into gentleMACS C Tubes using a Miltenyi OctoDissociator (predefined programme: 37C-h-TDK_1) for 1 hour at 37 °C.

Dissociated cells were filtered through a 100 µm cell strainer (Appleton Woods), washed with DPBS, and centrifuged at 800 x g for 4 min at room temperature to remove digestion buffer and cellular debris. The cell pellet was resuspended in 1–5 mL ACK Lysing Buffer (Gibco) and incubated for 5 min at room temperature, followed by centrifugation at 800 x g for 4 min at room temperature. The cell pellet was assessed according to size and embedded in 100 % Matrigel (Corning) using either a 6-well plate format (Helena Biosciences) (7 × 40 µL droplets per well) or, for smaller pellets, a 12-well plate format (3 × 25 µL droplets per well). Matrigel containing epithelial organoids was incubated for 20 min at 37 °C to allow polymerisation. Organoid Expansion medium supplemented with Y-27632 ([final 10 µM]; (Sigma-Aldrich)) was then added. Medium supplemented with Y-27632 was maintained during the initial culture period (4 days), after which fresh medium was replaced every 2 days (without Y-27632) until organoids reached confluency (approximately day 7). Organoids were subsequently passaged, expanded, and maintained until passage 5 (P5), at which point they were cryopreserved in liquid nitrogen and later subjected to a freeze–thaw recovery cycle test.

#### CAF Ligand Validation

Four PDO lines were cultured as described above and, on day 6, either co-cultured with CAFs or treated with all possible combinations (full factorial array) of 50 ng/mL HGF (Peprotech 100-39H-100UG), 500 ng/mL IL1R1, 50 ng/mL GREM1 (Peprotech 120-42-100UG) and 1 nM PGE_2_ (Biotechne 2296/10) prepared in base media, treatments were refreshed every 24 hours for 72 hours. The cultures were then analysed by TOB*is* MC.

#### Generation of Stable *PTGS2* CRISPR Knockout CRC CAFs

Anti-*PTGS2* Edit-R crRNA sequence:TGTACCCGGACAGGATTCTA (Horizon CM-004557-02-0002) and Edit-R tracrRNA was reconstituted to 50 µM with 10 mM Tris and duplexed by combining at a 1:1 ratio and incubating at room temperature for 30 min. Then 2.75 µL of 50 µM duplexed gRNA was mixed with 4.4 µL Lullaby transfection reagent and 220 µL OptiMEM and incubated for 20 min before being transferred into one well of a 12-well multi-well plate with 0.44×10^4^ Cas9 CAFs suspended in 880 µL 2% FBS DMEM. The Cas9 CAFs were maintained at 5% CO2, 37°C for 96 hours before harvesting and diluting to 1 cell per 100 µL and dispensing in 96-well multi-well plate. Colonies were selected, expanded and assessed for knockout. DNA was extracted from the clones using Quick-DNA Microprep Plus Kit (Zymo Research D7005T) and submitted for Sanger sequencing. Sequences were analysed using DECODR (https://www.decodrinc.com) [70]. Clones were expanded and embedded in Matrigel as previously described. Media was harvested from these cultures and PGE_2_ concentration assessed via ELISA (antibodies.com A78615).

#### PGE_2_ Rescue and Aspirin Treatments

Four PDO lines were cultured as described above and on day 6, for the rescue experiment, were co-cultured with unedited CAF or *PTGS2* KO CAF and treated with 0-50 nM PGE_2_. For the Aspirin study, PDO was cultured +/-CAFs, treated with 0.5 mM Aspirin (Bio-Techne 4092/50) or DMSO and treated +/-50 nM PGE_2_. All treatments were made up in base media and refreshed every 24 hours for 72 hours. The cultures were then processed by TOB*is* MC.

#### PGE_2_ Signalling Analysis

PDO were prepared as described above and after 24h the PDO cultures were treated with 50 nM PGE_2_. The cultures were then fixed in 4% PFA/PBS for TOB*is* MC at 0.5, 1, 2, 6 and 24 hours. The remaining cultures had base media change, without PGE_2_ supplementation, every 24 hours and were fixed for TOB*is* MC 48, 72, 96, 120 hours post PGE_2_ treatment.

#### Xenium FFPE *in situ* Protocol

Xenium *in situ* slides were prepared in accordance with the manufacturer’s instructions. Formalin-fixed paraffin-embedded (FFPE) colorectal tumour tissue blocks (Table S2) were sectioned at 5 µm thickness and mounted onto Xenium slides (capture area: 2.35 cm^2^). Sections were air-dried at room temperature for 30 min, followed by incubation at 42 °C for 3 hours. Tissues were subsequently deparaffinised and subjected to de-crosslinking to enable probe hybridisation. Xenium Prime probes (5,001 genes), together with a custom panel comprising an additional 111 gene targets (see Supplementary File ‘Custom_Xenium_Gene_Probes.csv’), were hybridised to the tissue overnight. This was followed by probe ligation and annealing of rolling circle amplification primers. Wash steps were performed to minimise autofluorescence. Nuclear staining with DAPI and application of segmentation reagents (interior RNA, interior protein and boundary markers) were carried out according to the Xenium user guide. Imaging was performed using the automated Xenium Analyzer, which quantified transcripts via gene-specific optical barcodes. Cell segmentation was determined using integrated signals from DAPI, interior and boundary staining channels.

#### Xenium Single-cell Transcriptomics: Annotation

Xenium-derived gene expression matrices were imported into R and analysed using Seurat [71]. Raw count matrices were used to generate Seurat objects, and cells with zero or invalid total counts were excluded. Quality control filtering retained cells with a minimum of 50 detected features and 100 total transcript counts. Data were log-normalised, and the 2,000 most variable features were identified for downstream analysis. Scaled data were subjected to principal component analysis (PCA), followed by construction of a nearest-neighbour graph. Uniform Manifold Approximation and Projection (UMAP) was used for dimensionality reduction and visualisation, and graph-based clustering was performed to define cell populations. Cluster-specific marker genes were identified by differential expression analysis, and cluster identities were assigned based on canonical marker gene expression profiles. This enabled the delineation of epithelial cell populations and *PTGS2*^*+*^ fibroblast subsets. Xenium 5K proCSC and revCSC gene signature can be found in Supplementary File ‘CRC_Xenium_Signatures.csv’.

#### Xenium Single-cell Transcriptomics: Spatial Analysis

All spatial analyses were performed in Python using MuSpAn (v1.2.4) [22]. Raw Xenium outputs were converted into MuSpAn domain objects, in which each cell was represented as a point object defined by the centroid of the corresponding 10X Genomics segmentation mask and annotated with cell-type labels and raw transcript counts (see Data Availability). The exterior boundary of each tissue domain was estimated using an alpha-shape reconstruction with a radius parameter of 200 µm [22]. To facilitate visualisation, quality control and macro–scale analysis, each spatial domain was further partitioned using a hexagonal lattice with a side length of 1,240 µm. Transcript counts for *PTGS2* and *DACH1* were assigned to the resulting hexagonal tiles for inspection of large-scale spatial structure and downstream validation.

##### Topographical Correlation Analysis

To quantify the local spatial relationship between *PTGS2*^+^ fibroblasts and *DACH1*^+^ epithelial cells, we applied the Topographical Correlation Map (TCM), a local indicator of spatial association implemented in MuSpAn [22]. Briefly, TCM evaluates the local density of a target population (population B) within a specified interaction radius, *r*, around each point in a source population (population A), and compares the observed density with that expected under complete spatial randomness. The observed-to-expected ratio is linearly transformed such that values of 1 indicate maximal co-localisation, −1 indicate maximal spatial exclusion, and 0 correspond to the null expectation. These local statistics are then spatially propagated using Gaussian kernels centred on each point in population A, with kernel variance *σ*, to generate a continuous topographical map of local spatial association. TCM was computed from *PTGS2*^+^ fibroblasts to *DACH1*^+^ epithelial cells using an interaction radius of 50 µm and a Gaussian kernel variance of 50.

##### Cross-pair Correlation Analysis

To assess spatial association across multiple length scales, we computed the cross-pair correlation function (cross-PCF) between *PTGS2*^+^ fibroblasts and *DACH1*^+^ epithelial cells, as well as between *PTGS2*^+^ fibroblasts and the full epithelial compartment [22]. The cross-PCF, *g*_*A,B*_(*r*), estimates the relative frequency of point pairs from populations A and B separated by distance *r* compared with the expectation under complete spatial randomness. Values of *g*_*A,B*_(*r*) *>* 1 indicate enrichment or co-localisation at distance *r*, whereas values of *g*_*A,B*_(*r*) *<* 1 indicate spatial exclusion.

To control for large-scale spatial heterogeneity, cross-PCF was calculated within hexagonal tiles containing more than 30 cells from each of the two populations under comparison. Calculations used an annulus width of 20 µm and a maximum distance of 600 µm, corresponding to less than half the diameter of each hexagonal region, with automatic boundary correction supported. Cross-PCF curves were then averaged across all eligible hexagonal regions, and 95% confidence intervals around the mean were calculated.

##### Nearest-neighbour Analysis

Nearest-neighbour distance distributions were computed as Euclidean distances from each *PTGS2*^+^ fibroblast to the nearest *DACH1*^+^ epithelial cell, and separately to the nearest cell within the full epithelial compartment. Analyses were restricted to hexagonal tiles containing more than 30 cells in each population to ensure sufficient spatial coverage. Distance distributions were computed for each tile and subsequently pooled across all samples to generate an aggregated nearest-neighbour distribution.

##### Distance-resolved epithelial profiling

To quantify how epithelial molecular features varied as a function of proximity to *PTGS2*^+^ fibroblasts, we constructed a spatial distance matrix between epithelial cells and *PTGS2*^+^ fibroblasts, defined as cells with at least 10 *PTGS2* transcripts per cell. For each epithelial cell, the Euclidean distance to the nearest *PTGS2*^+^ fibroblast was computed and binned in 50 µm intervals up to a maximum distance of 500 µm. These distance bins were used to evaluate transcript density and spatial gene-signature scores within the epithelial compartment as a function of fibroblast proximity.

##### Spatial Visualisation

Transcript density across tissue sections was visualised in Xenium Explorer (v4.0.0; 10x Genomics). Spatial gene signatures were defined using representative marker genes and co-visualised with the Gene Visualization Gallery in Xenium Explorer to generate composite maps of spatial expression.

### Statistical Analyses

All statistical analyses were performed using GraphPad Prism 11. *P* values of less than 0.05 were considered statistically significant.

## Supporting information

Secretome_CRISPR_Library

CRC_scRNA-seq_Modules_All

CRC_Xenium_Signatures

Custom_Xenium_Gene_Probes

## Data Availability

Raw SPLiT-Seq data are available via BioProject PR-JNA1308308 (https://www.ncbi.nlm.nih.gov/bioproject/1308308). Processed SPLiT-Seq data are available at Zenodo (https://doi.org/10.5281/zenodo.20599066). Raw and processed CyTOF data and illustrations are available as a Community Cytobank project 1675 (https://community.cytobank.org/cytobank/projects/1675). Pro-cessed Xenium data are available at Zenodo (https://doi.org/10.5281/zenodo.20446905 and https://doi.org/10.5281/zenodo.20511420).

## Acknowledgements

We are extremely grateful to M. Garnett, H. Francies and the Cell Model Network UK for sharing CRC PDOs and O. De Wever for sharing CRC CAFs. We thank Y. Guo and the UCL CI Flow-Core for CyTOF support. We acknowledge the UCL Organoid Translational Technology Platform for deriving and supplying the organoid models to use in this research and the UCLH Biobank for Studying Health and Disease for providing the human tissue samples and the clinical data that was used in and supported the generation of the organoid models. This work was supported by the Constance Travis Trust, the Biotechnology and Biological Sciences Research Council (BB/T008709/1), UKRI Medical Research Council (MR/T028270/1), Cancer Research UK (C60693/A23783), the Cancer Research UK City of London Centre (CTRQQR-2021/100004), and the UCLH Biomedical Research Centre (BRC422). SJL was supported by CRUK Programme Grant (DRCNPG-Jun22/100002). Mathematical and biological integration was supported by CRUK CRC-STARS Strategic Grant (SEBCRCS-2024/100001). JWM was supported by CRUK (CTRQQR-2021\100002), through the Cancer Research UK Oxford Centre. E.J.M. was supported by the Lee Placito Research Fellowship (University of Oxford).

## Author Contributions

C.M. performed all PDO+CAF CRISPR screening experiments, ligand validation, and analysis. R.O. performed all PDO+CAF SPLiT-seq experiments and analysis. E.J.M. and J.W.M. performed all Xenium analysis. N.L. performed all PDO chemosensitivity experiments and analysis. P.V. established CRC PDOs. R.A. performed computational analysis of CRC patient data. A.D. prepared ligand and drug arrays. S.C. and J.L performed antibody conjugations. A.W. performed Geodesic Sinkhorn analysis. M.L.A produced CAF-conditioned media. A.C. established Cas9 CAFs. J.C. constructed VR landscapes. S.K. supervised Geodesic Sinkhorn analysis. P.D. supervised computational analysis of CRC patient data. S.L. supervised Xenium analysis. C.J.T. conceived the project, designed the study, analysed the data, and wrote the paper.

## Supplementary Information

**Table 1.** Antibodies used for TOB*is* MC experiments.

| Extracellular Antigen | Metal | Clone | Supplier | Antibody Registry |
| --- | --- | --- | --- | --- |
| CD276 (B7-H3) | <sup>112</sup>Cd | B7C23 | Prof. Kerry Chester | — |
| FAP | <sup>112</sup>Cd | Polyclonal | R&D Systems | AB_2102369 |
| CD326 (EpCAM) | <sup>113</sup>In | 9C4 | BioLegend | AB_756075 |
| CHGA | <sup>141</sup>Pr | C-12 | Santa Cruz | AB_2801371 |
| L1CAM | <sup>146</sup>Nd | L1-OV198.5 | BioLegend | AB_2616786 |
| FAP | <sup>152</sup>Sm | Polyclonal | R&D Systems | AB_2102369 |
| TACSTD2 (TROP2) | <sup>167</sup>Er | NY18 | BioLegend | AB_2564376 |
| EPHB2 | <sup>169</sup>Tm | 2H9 | BD Biosciences | AB_2738897 |
| CD55 | <sup>171</sup>Yb | JS11 | BioLegend | AB_314859 |
| Intracellular Antigen | Metal | Clone | Supplier | Antibody Registry |
| pHistone H3 [S28] | <sup>89</sup>Y | HTA28 | BioLegend | AB_1227660 |
| Vimentin | <sup>111</sup>Cd | RV202 | BD Biosciences | AB_393716 |
| Cytokeratin 18 | <sup>114</sup>Cd | C-04 | Abcam | AB_305647 |
| Pan-CK | <sup>115</sup>In | AE1/AE3 | BioLegend | AB_2616960 |
| GFP | <sup>116</sup>Cd | 5F12.4 | eBiosciences | — |
| cCaspase 3 [D175] | <sup>142</sup>Nd | D3E9 | CST | AB_10897512 |
| pMKK4/SEK1 [S257] | <sup>143</sup>Nd | C36C11 | CST | AB_2140946 |
| SOX2 | <sup>144</sup>Nd | Polyclonal | Abcam | AB_2341193 |
| pNDRG1 [T346] | <sup>145</sup>Nd | D98G11 | CST | AB_10693451 |
| OPTN | <sup>147</sup>Sm | E4P8C | CST | AB_3073769 |
| CDK1 | <sup>148</sup>Nd | A17 | Thermo | AB_2533105 |
| p4E-BP1 [T37/46] | <sup>149</sup>Sm | 236B4 | CST | AB_560835 |
| pRB [S807/811] | <sup>150</sup>Nd | J1112-906 | BD Biosciences | AB_647295 |
| pBAD [S112] | <sup>151</sup>Eu | 40A9 | CST | AB_560884 |
| pERK [T202/Y204] | <sup>152</sup>Sm | 20A | BD Biosciences | AB_399648 |
| ANXA1 | <sup>153</sup>Eu | D5V2T | CST | AB_2799031 |
| pSMAD1/5/9 | <sup>154</sup>Sm | D5B10 | CST | AB_2493181 |
| pAKT [S473] | <sup>155</sup>Gd | M89-61 | BD Biosciences | AB_2737620 |
| pNF-κB [S529] | <sup>156</sup>Gd | K10-895.12.50 | BD Biosciences | AB_647284 |
| Isotype Control | <sup>157</sup>Gd | MOPC21 | Abcam | AB_2736846 |
| pP38 [T180/Y182] | <sup>158</sup>Gd | D3F9 | CST | AB_2139682 |
| Cas9 | <sup>159</sup>Tb | 7A9-3A3 | CST | AB_2750916 |
| KI67 | <sup>160</sup>Gd | D3B5 | CST | AB_2687446 |
| pLATS1/2 [S872/S909] | <sup>161</sup>Dy | EPR27262-6 | Abcam | — |
| pC-JUN [S73] | <sup>162</sup>Dy | D47G9 | CST | AB_2129575 |
| p21 Waf1/Cip1 | <sup>163</sup>Dy | 12D1 | CST | AB_3099451 |
| TOP2A | <sup>164</sup>Dy | D10G9 | CST | AB_2797871 |
| DACH1 | <sup>165</sup>Ho | 3B6D2 | Proteintech | AB_2089711 |
| pGSK-3β [S9] | <sup>166</sup>Er | D85E12 | CST | AB_10013750 |
| pSMAD2/3 | <sup>168</sup>Er | D27F4 | CST | AB_2631089 |
| pMEK1/2 [S221] | <sup>170</sup>Er | 166F8 | CST | AB_490903 |
| BIRC3 | <sup>172</sup>Yb | OTI3G4 | Thermo | — |
| pS6 [S240/S244] | <sup>173</sup>Yb | D68F8 | CST | AB_10694233 |
| cPARP [D214] | <sup>174</sup>Yb | D64E10 | CST | AB_10699459 |
| pC-FOS [S32] | <sup>175</sup>Yb | D82C12 | CST | AB_10557109 |
| Cyclin B1 | <sup>176</sup>Yb | GNS-11 | BD Biosciences | AB_395290 |

**Table 2.**
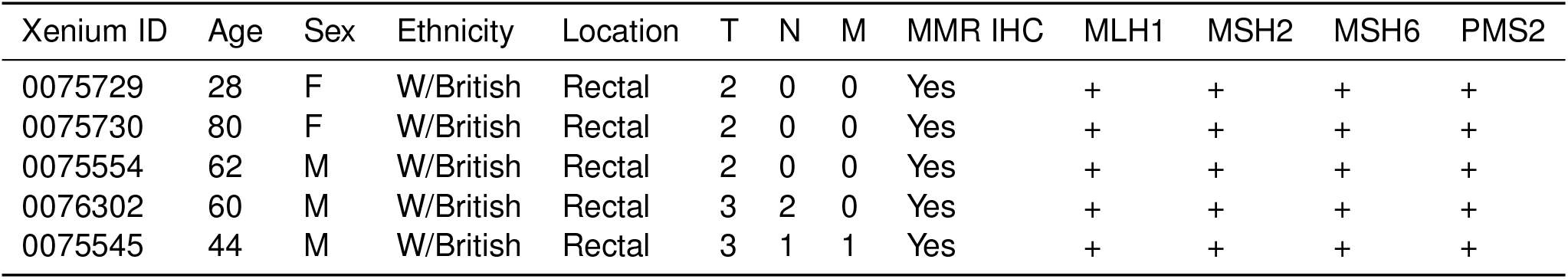
Xenium 5K Patient Metadata.

### Supplementary Figures

**Figure S1.**
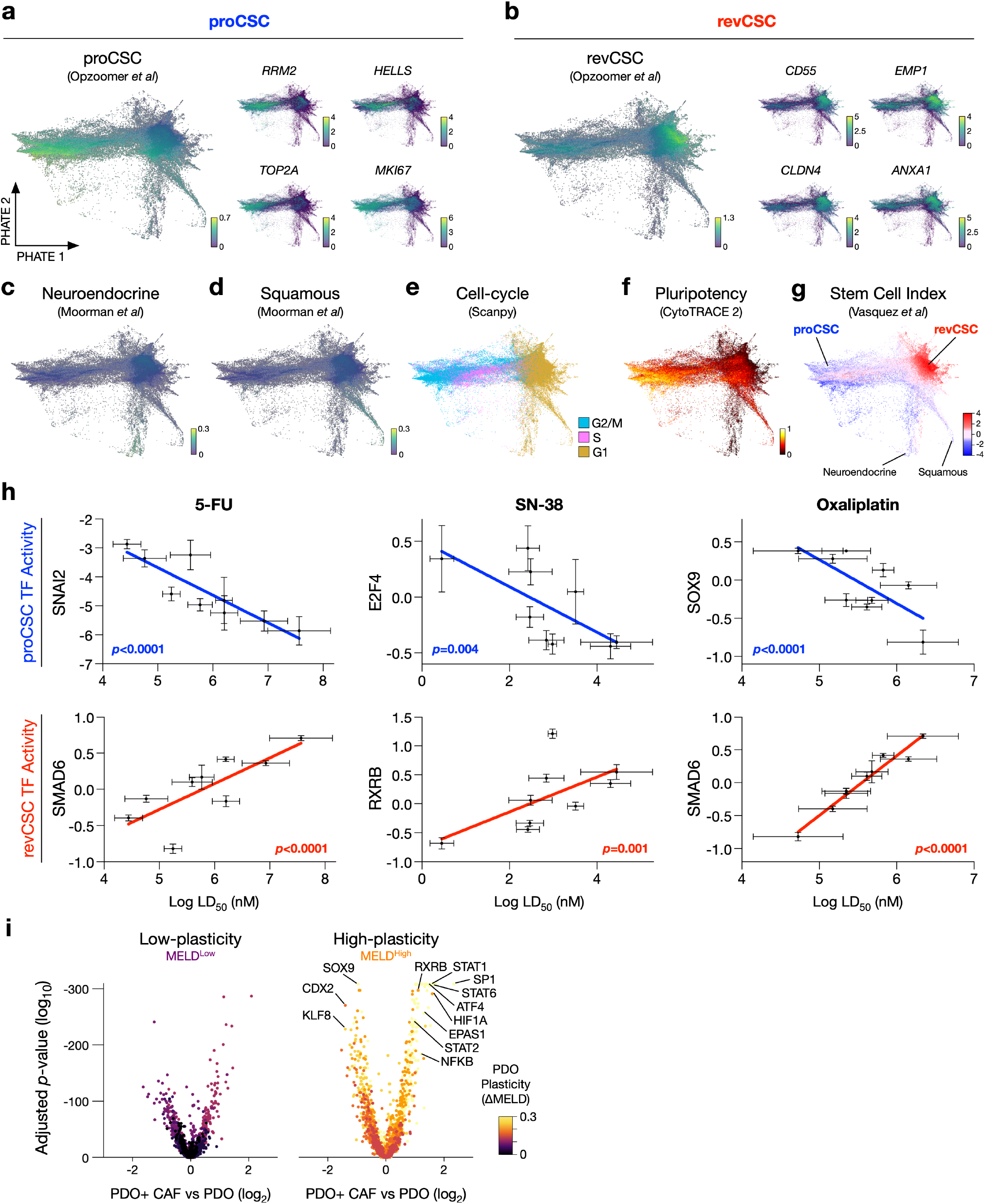
Single-cell RNA-seq Analysis of CRC PDOs +/-CAFs. **a)** scRNA-seq PHATE of x10 CRC PDOs (89,894 single cells) +/-CAFs (in triplicate), coloured by proCSC gene signature and canonical genes, **b)** revCSC gene signature and canonical genes, **c)** neuroendocrine gene signature, **d)** squamous gene signature, **e)** cell-cycle phase, **f)** predicted pluripotency, and **g)** Stem Cell Index (z-score). **h)** PDO proCSC and revCSC transcription factor activities versus 5-FU, SN-38 (Irinotecan), and Oxaliplatin LD_50_. Pearson correlation. **i)** Transcription factor activities +/-CAFs in low-plasticity MELD^Low^ and high-plasticity MELD^High^ PDOs.

**Figure S2.**
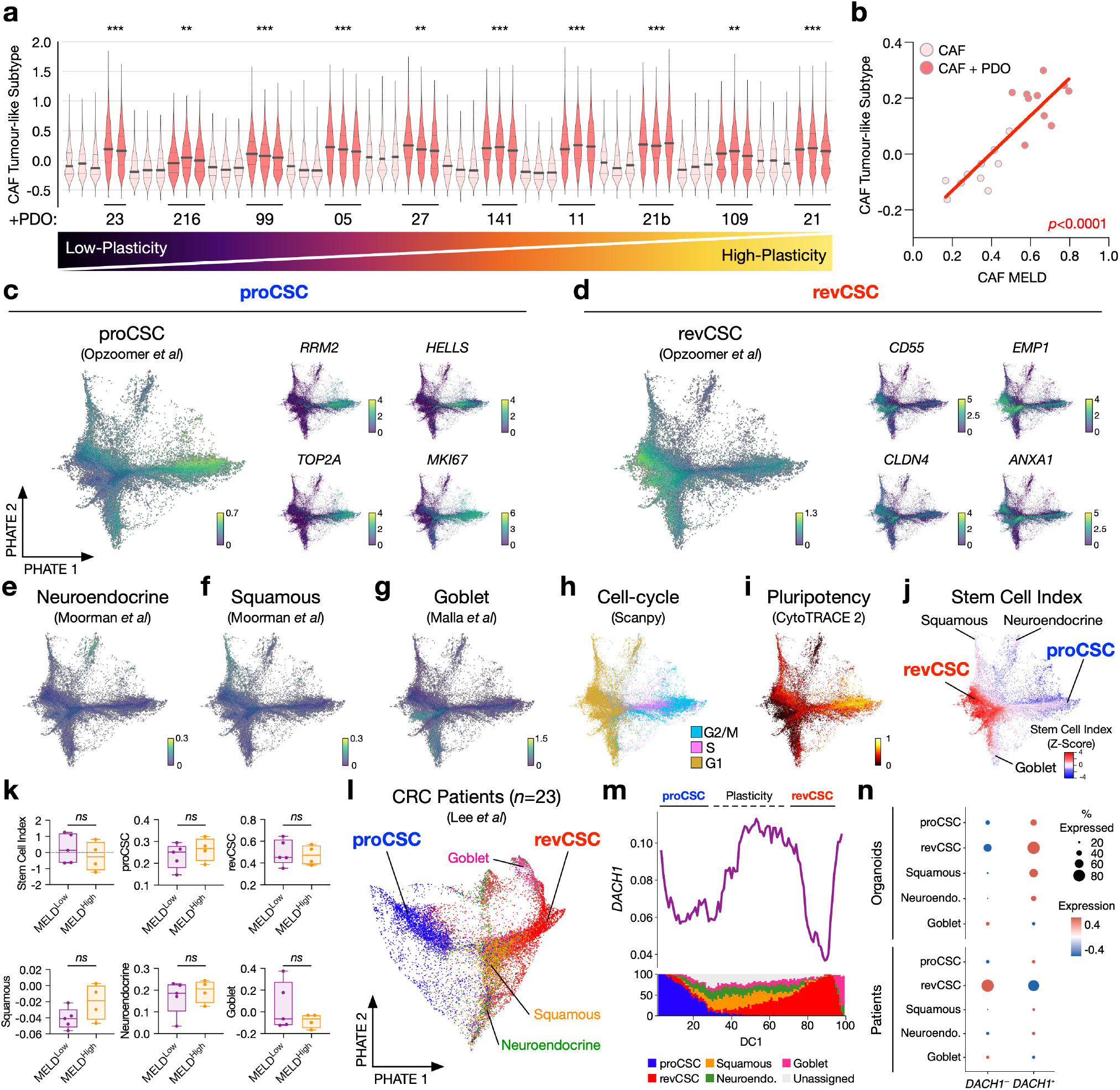
scRNA-seq Characterisation of CRC CAFs and Epithelial Cells. **a)** CAF Tumour-like gene expression signature for each CRC CAF +/-CRC PDOs ranked by PDO intrinsic-to-extrinsic fate. Unpaired *t*-test of means per CAF culture. **b)** Mean CAF Tumour-like gene expression signature versus mean CAF MELD per PDO. Pearson correlation. **c)** scRNA-seq PHATE of x10 CRC PDO monocultures (in triplicate) (39,022 single cells), coloured by proCSC gene signature and canonical genes, **d)** revCSC gene signature and canonical genes, **e)** neuroendocrine gene signature, **f)** squamous gene signature, **g)** goblet gene signature, **h)** cell-cycle phase, **i)** predicted pluripotency, and **j)** proCSC-to-revCSC stem cell index. **k)** Stem cell index, proCSC, revCSC, squamous, neuroendocrine, and goblet gene signatures in MELD^Low^ and MELD^High^ PDOs. Welch’s t-test, ns = not significant. **l)** Single-cell PHATE of human CRC epithelia (SMC Cohort [18]) coloured by gene expression signatures (17,469 cells). **m)** *DACH1* expression and cellular composition ordered by diffusion component (DC) 1. **n)** Expression of *DACH1* in proCSC, revCSC, Neuroendocrine, Squamous, and Goblet cells in organoids and patients.

**Figure S3.**
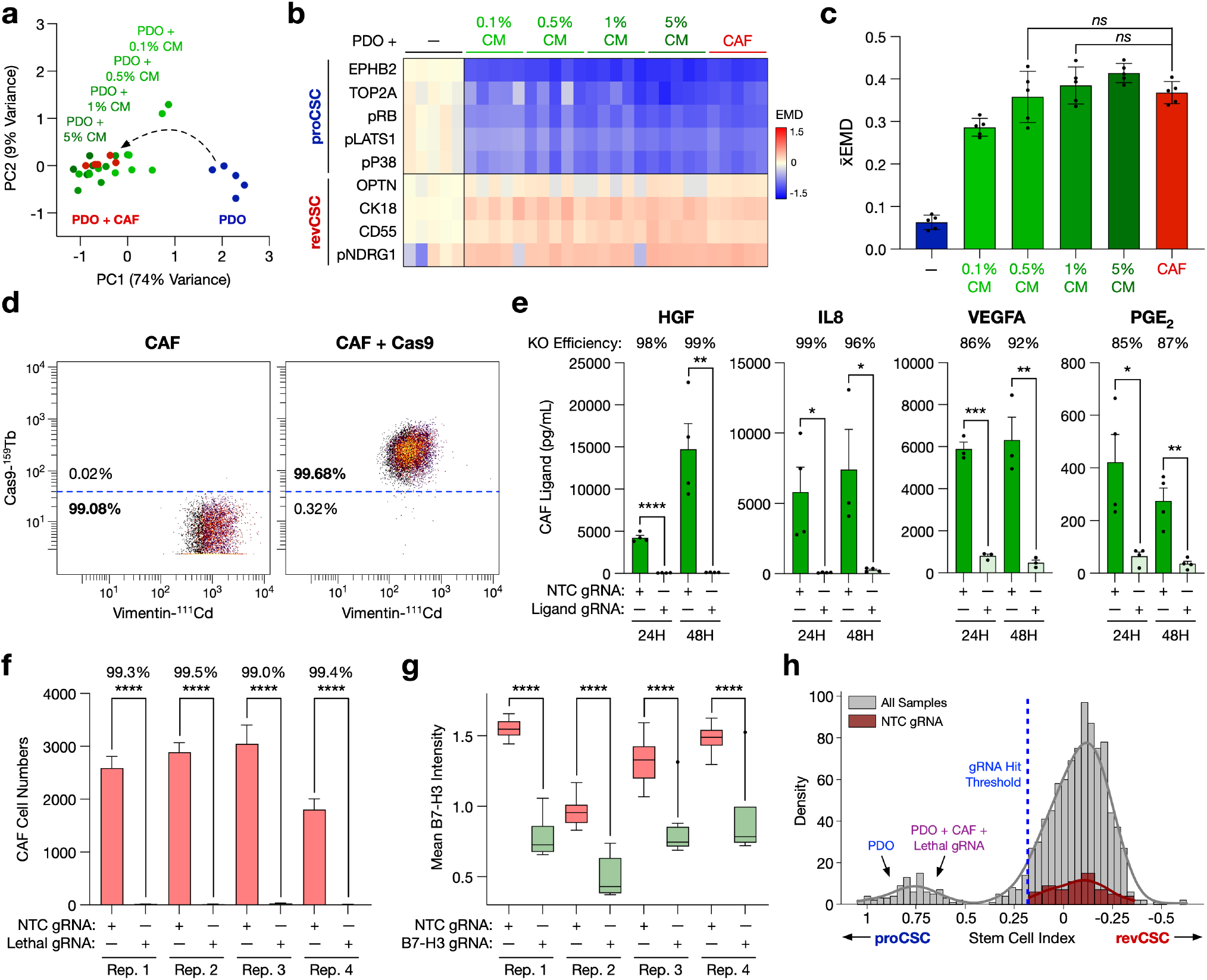
Single-cell CRISPR Screening of CAF-PDO Intercellular Signalling. **a)** TOB*is* MC EMD-PCA, **b)** EMD heatmap, and **c)** *x*EMD analysis of PDOs grown alone, in co-culture with CAFs, or treated with 0.1%, 0.5%, 1%, and 5% CAF conditioned media (CM) (*n* = 5). Unpaired *t*-test. **d)** Single-cell CRC CAF Cas9 expression. **e)** Quantitative CRISPR knock out (KO) efficiency of native HGF, IL8, VEGFA, and PGE_2_ from CRC CAFs compared to non-targeting control (NTC) guide RNA (gRNA). **f)** CAF cell numbers throughout the PDO-CAF CRISPR screen following transfection with NTC or lethal gRNA. Unpaired *t*-test. **g)** B7-H3 mean intensity following transfection with NTC or B7-H3 gRNA. Unpaired *t*-test. **h)** PDO Stem cell index histogram of all samples and NTC gRNA. Paired *t*-test. **** = *p* < 0.0001, *** = *p* < 0.001, ** = *p* < 0.01, * = *p* < 0.05, ns= not significant.

**Figure S4.**
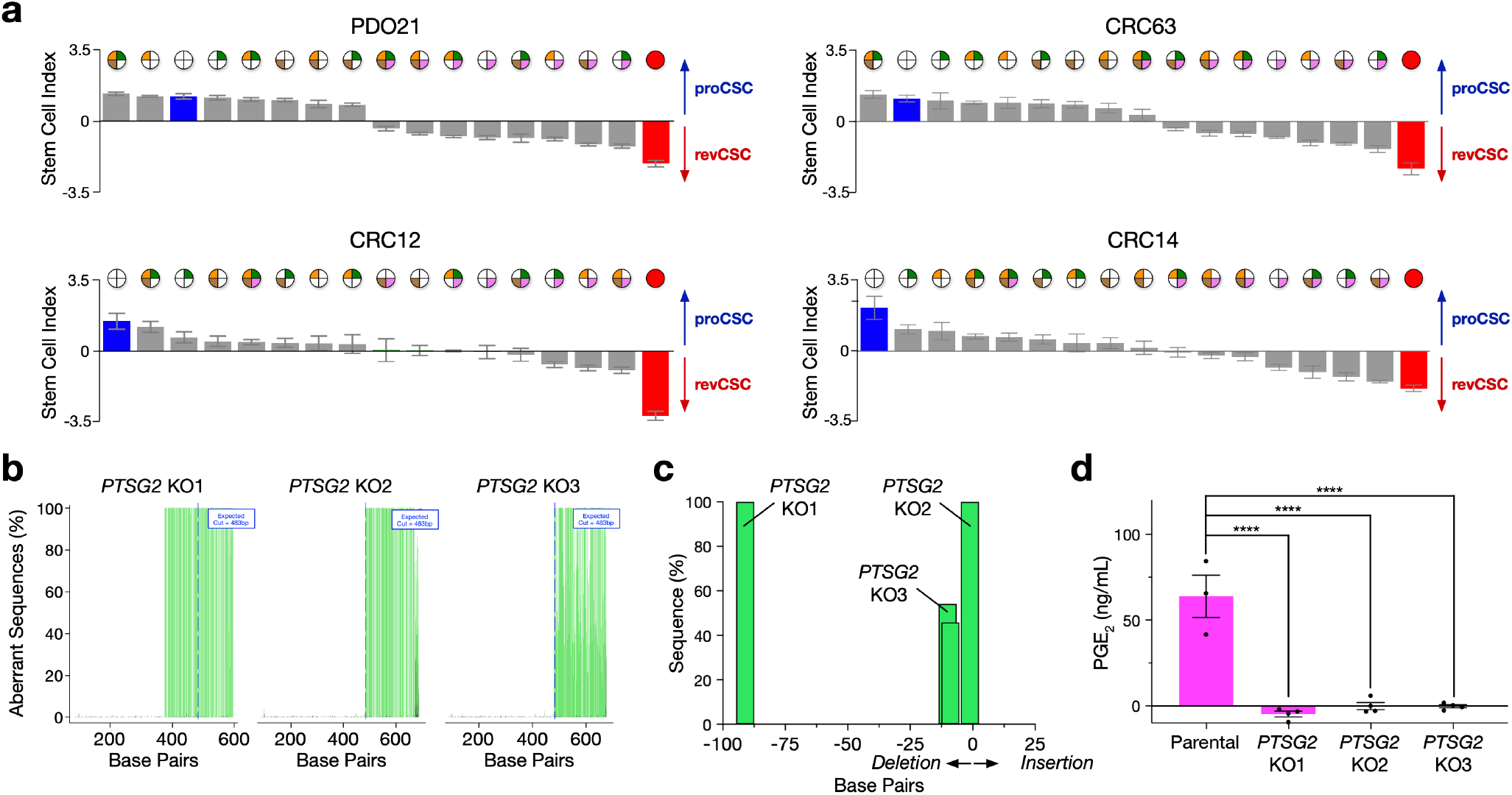
CAF Ligand Validation. **a)** Stem cell index for each ligand combination per PDO. **b)** and **c)** TIDE analysis of CAF *PTGS2* CRISPR KO clones. **d)** PGE_2_ ELISA of CAF conditioned media from stable CAF *PTGS2* CRISPR KO clones.

**Figure S5.**
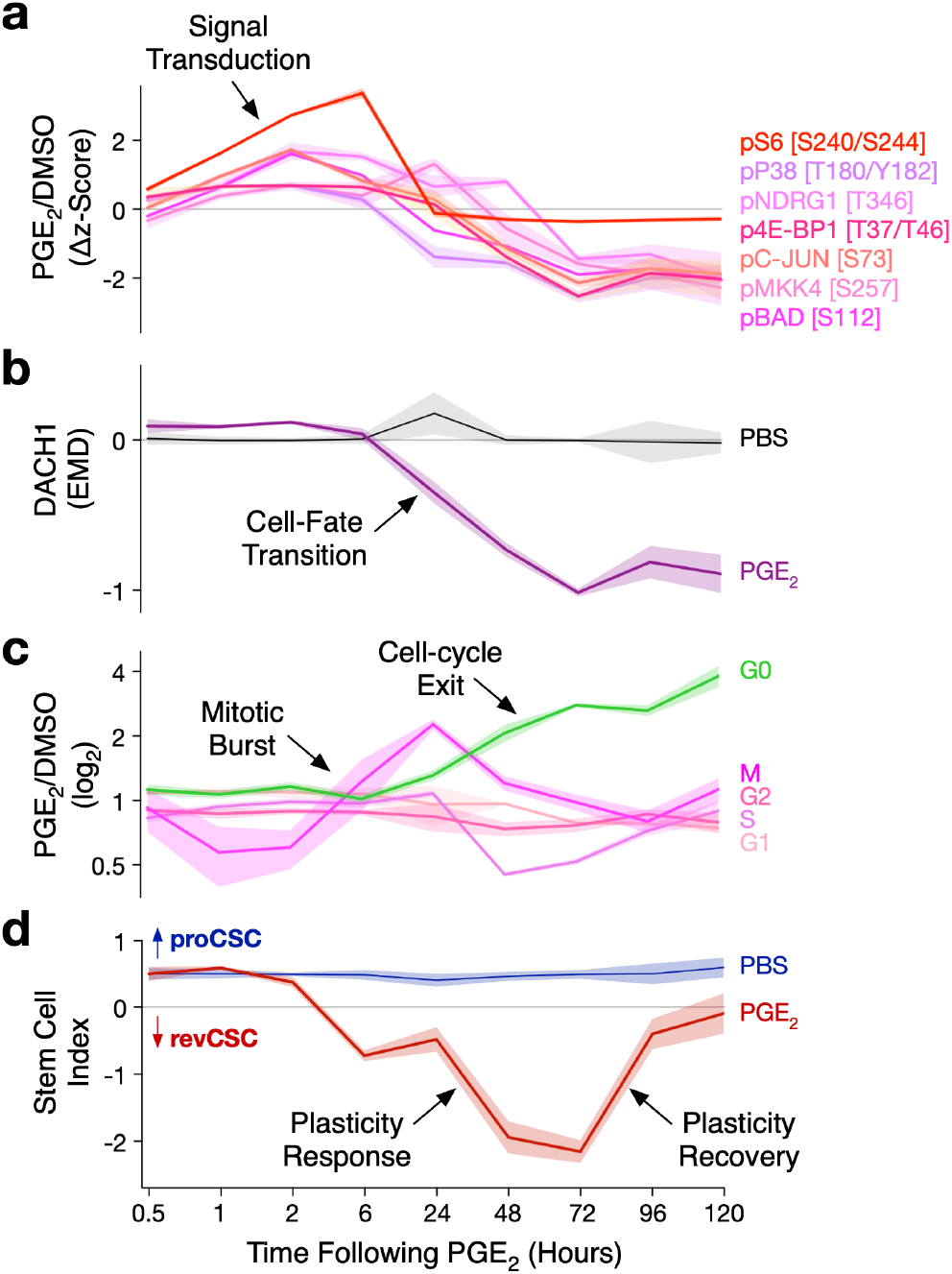
PGE_2_ Signalling Dynamics. **a)** Single-cell TOB*is* MC analysis of CRC PDOs treated +/-50 nM PGE_2_ analysing epithelial phospho-signalling, **b)** DACH1, **c)** cell-cycle, and **d)** stem cell index.

**Figure S6.**
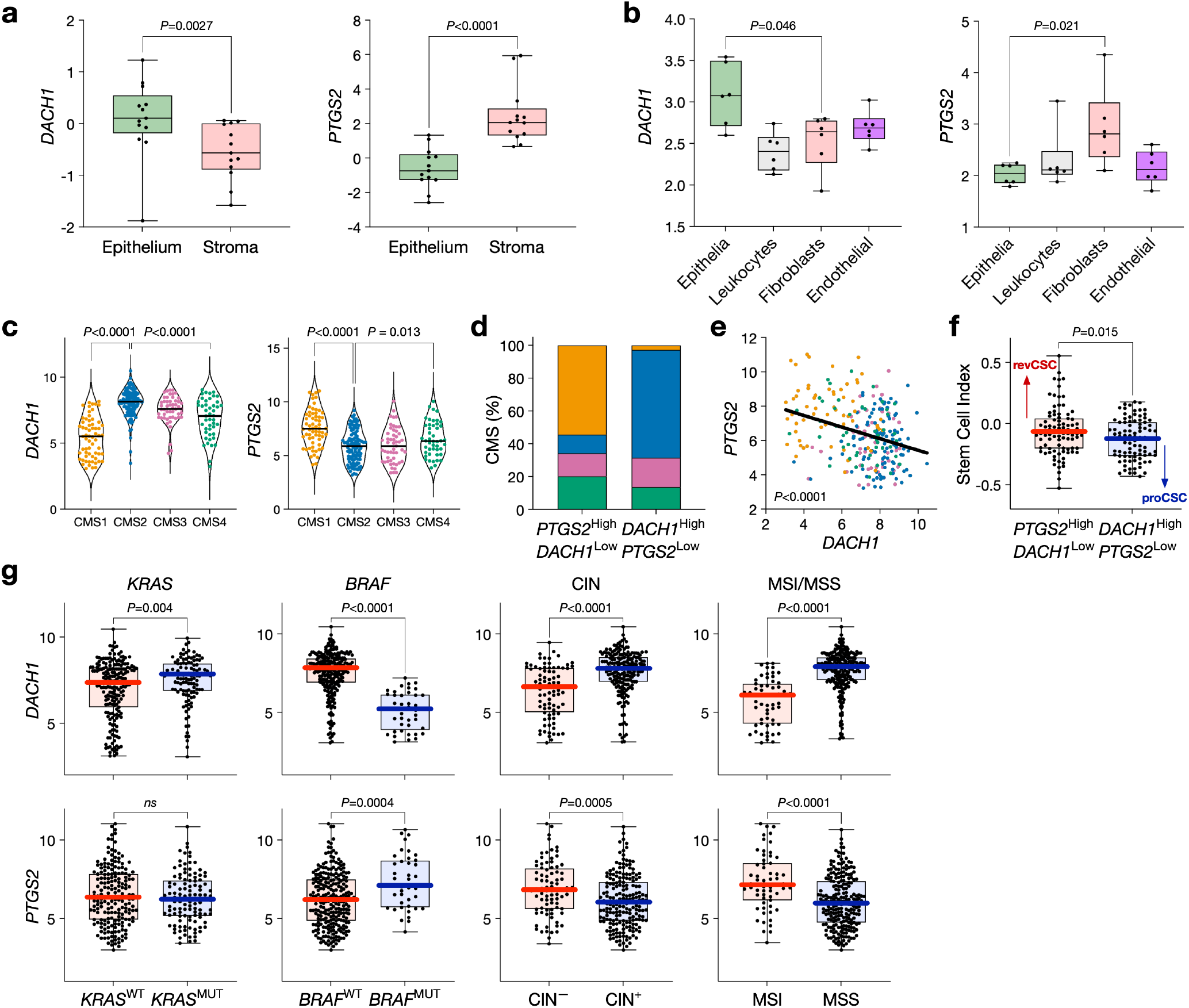
*DACH1* and *PTGS2* in Human CRC. **a)** *DACH1* and *PTGS2* expression in epithelial and stromal laser capture dissected human CRC. **b)** *DACH1* and *PTGS2* expression in FACS sorted epithelia, leukocytes, fibroblasts, and endothelial cells. **c)** *DACH1* and *PTGS2* expression across CRC Consensus Molecular Subtypes (CMS). **d)** Relative proportion of CMS subtypes in *PTGS2*^*High*^/*DACH1*^*Low*^ and *PTGS2*^Low^/*DACH1*^*High*^ CRC tumours. **e)** Anti-correlation between *DACH1* and *PTGS2* expression in human CRC. Pearson correlation. **f)** Stem cell index of *PTGS2*^*High*^/*DACH1*^*Low*^ and *PTGS2*^Low^/*DACH1*^*High*^ CRC tumours. **g)** *DACH1* and *PTGS2* expression across KRAS, BRAF, CIN, and MSS/MSI status in human CRC.

**Figure S7.**
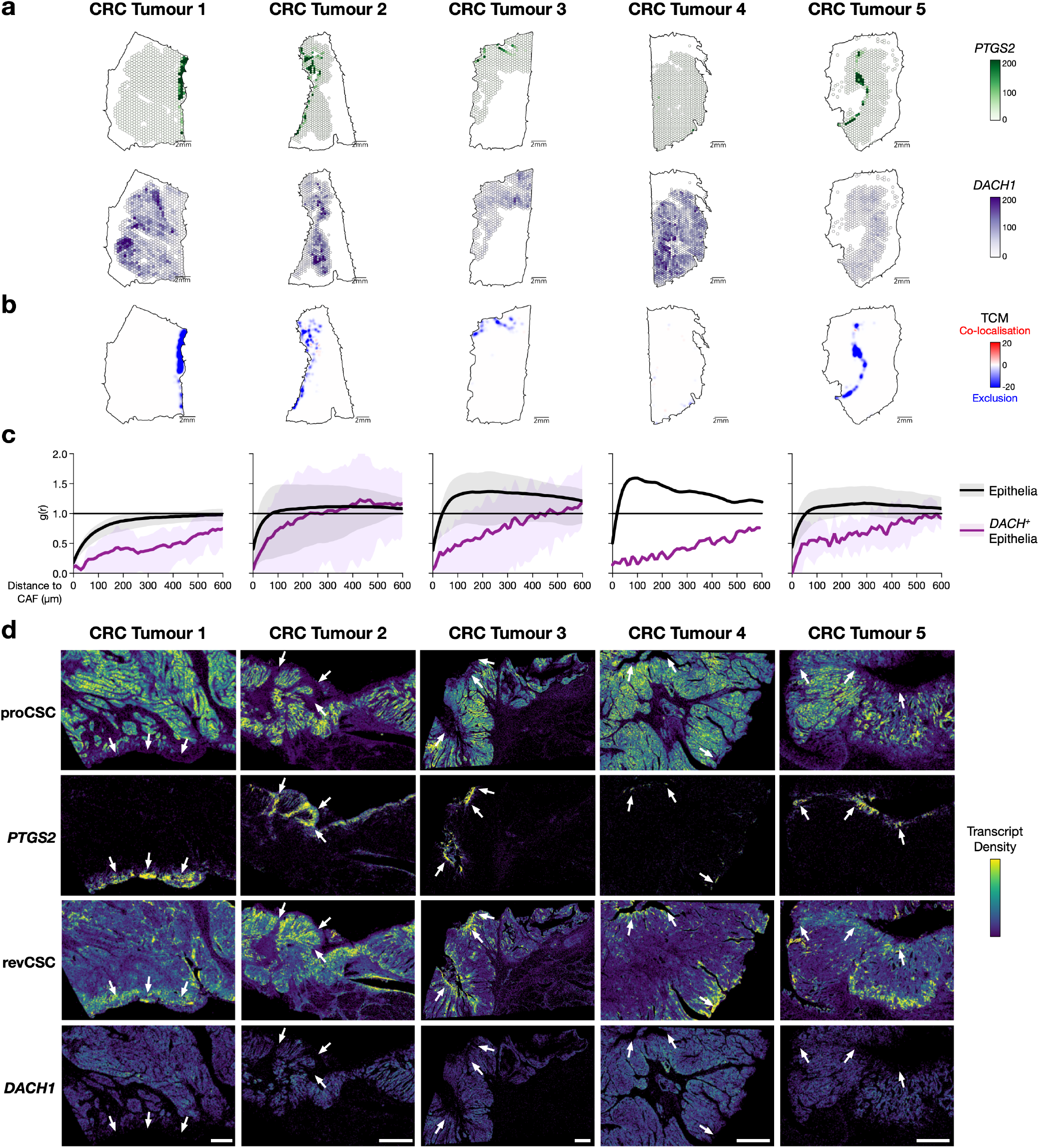
Xenium 5K Analysis of Primary Human CRC. **a)** CAF *PTGS2* and epithelial *DACH1* expression of 5 human primary CRC tumours analysed by Xenium 5K (hexgrid side length 200 µm). **b)** Topographical Correlation Map of *PTGS2*:*DACH1* co-localisation and exclusion (50 µm radius). **c)** Cross-pair correlation function analysis of all CAFs and all epithelia (black) versus *PTGS2*^*+*^ CAFs and *DACH1*^*+*^ epithelia (purple). Mean, 95% CI. **d)** Transcript densities of *PTGS2* and proCSC and revCSC gene signatures, and *DACH1* in human CRC tumours. Arrows indicate areas of *PTGS2* expression.

